# Ecological Cascades and Future Hantavirus Spillover Risk in a Changing Climate

**DOI:** 10.64898/2026.07.13.738318

**Authors:** Pranav S. Pandit, Andie H. Jian, Ayushi Gupta, Pranav S. Kulkarni

**Author notes:** These authors contributed equally to the manuscript (Pranav S. Pandit, Pranav S. Kulkarni).

## Abstract

Anthropogenic climate change is reshuffling global biodiversity and accelerating zoonotic spillover at the human-wildlife interface. The 2026 multi-country cruise ship outbreak of Andes virus highlights the unpredictable and globalized threat posed by rodent-borne orthohantaviruses, which cause severe, highly fatal human diseases worldwide. Despite this significant public health risk, macroecological synthesis linking environmental drivers to systemic shifts in hantavirus hazards has been lacking. Here, we implement an integrated ecological and machine learning framework combining 89 unique virus-host associations across 61 reservoir species with mechanistic Force-of-Infection (FOI) modeling to project global spillover hazards under current and future climate scenarios (SSP2-4.5 and SSP5-8.5) for 2041–2060. Our models demonstrate a spatially uneven but geographically extensive expansion of spillover hazards, with approximately 74.9% of modeled land area experiencing an increase in FOI. We identify three distinct high-hazard belts heavily concentrated in already-temperate reservoir ranges: western and central North America, central and northeastern Europe extending into western Russia, and discrete patches within central South America. Crucially, our findings reveal that hantavirus hazards are fundamentally shaped along virus-specific ecological axes. While thermal and precipitation anomalies predominate the transmission dynamics of Old and New World lineages like Sin-Nombre virus and Thailand virus, distinct urban-centric Profiles emerge for Seoul virus and Mamore virus, driven explicitly by fine-scale anthropogenic landscape modifications. These results reveal that climate change will not simply intensify viral spillover uniformly but will fundamentally restructure it across varied climatic and human-dominated environments. Our maps provide a critical baseline for targeted global surveillance and highlight the urgent need to integrate spatial conservation planning with public health preparedness.

## Introduction

The 2026 multi-country outbreak of Andes virus (ANDV), identified on a cruise ship and involving twelve laboratory-confirmed (one unconfirmed) cases of hantavirus-associated respiratory illness, has highlighted an urgent need for international surveillance efforts to assess ongoing risks of exposure to rodent-borne hantaviruses and to evaluate their potential for further spread globally^1^. Hantaviruses are single-stranded RNA viruses (family Bunyaviridae, genus *Orthohantavirus*) and can cause pulmonary syndromes and hemorrhagic fevers with high case fatality rates.

Orthohantaviruses pose public health risk globally, and it is estimated that somewhere between 10,000 and 200,000 infections occur annually worldwide^2^. Asia, where Hantaan (HTNV) and Seoul viruses (SEOV) cause hemorrhagic and renal syndrome affect tens of thousands annually with case fatality rates reaching up to 5-15%. Europe reports several thousand cases of Puumala virus (PUUV) and Dobrava-Belgrade virus (DOBV). SEOV represents a global threat due to its association with *Rattus* species with distribution across Europe, Asia, Africa, and North America. Severe cases of Sin Nombre virus (SNV) causes hantavirus pulmonary syndrome in North America with 30-40% case fatality, while ANDV with 50% cases fatality is reported to have shown limited human to human transmission post initial spillover in South America. Orthohantaviruses in Africa are relatively less characterized with reports of Tanganya virus (TANGV) in Tanzania, Sangassou virus in Guinea (SANGV), and SEOV in multiple sub-Saharan countries; suggesting broader geographic distribution than historically perceived.

Spillover events involving orthohantaviruses are epidemiologically linked to direct and close contact with infectious rodent reservoirs, primarily through exposure to contaminated rodent excreta, inhalation of aerosolized viral particles, and, less commonly, rodent bites. The risk of zoonotic transmission is strongly modulated by the dynamics of rodent reservoir populations and interactions of humans at specific human-animal interfaces that lead to high probability of epidemiological contacts. The distribution and abundance of rodent reservoir populations can be predicted by combinations of precipitation, temperature, vegetation cover, and landscape structure. These same variables are shifting across the globe under anthropogenic climate change^3^. Previous work has established that climate change will substantially alter the distributions of rodent reservoirs and their associated viruses. Our recent studies on rodent-borne arenaviruses (family *Arenaviridae*), which share similar transmission ecology with hantaviruses, demonstrated that projected climate shifts could significantly expand or contract endemic areas, directly impacting human spillover risk in South America^4,5^. However, specific climatic and landscape drivers of orthohantaviruses spillover risks are unknown at macroecological scale along with continental-scale projections.

We present an integrated ecological framework linking environmental and landscape drivers to hantavirus spillover hazard across all known zoonotic orthohantaviruses, building on our previously validated methodology applied to arenaviruses^5^. We integrate virus–host association records, rodent reservoir occurrence and sero-prevalence data, climatic and habitat variables that constrain reservoir distributions, and climate projections to estimate current and future hazard. We estimated a spillover hazard by quantifying the force of infection FOI, defined as the probability of successful transmission to humans, based on human-rodent interactions under two different climate change scenarios. In addition to it, these predictive models provide insight into the environmental drivers underlying changes in spillover hazard, revealing how factors such as landscape structure and shifts in precipitation patterns and seasonality shape future transmission potential. We focus on 12 orthohantaviruses with well-documented spillover events and broad geographic ranges, including the currently ongoing ANDV outbreak in South America.

## Results

### Diversity and distribution of orthohantavirus reservoirs

Among the 12 orthohantavirus species included in this analysis, we identified 61 unique vertebrate host species comprising 89 unique virus-host associations. Host records were overwhelmingly dominated by rodents of the families Cricetidae and Muridae (particularly Subfamilies Arvicolinae, Sigmodontinae, and Murinae), with the remaining belonging to the families Geomyidae, Gliridae, Heteromyidae, or Sciuridae. Through literature search, we identified the distribution of prevalences and seroprevalences of the 12 orthohantavirus by host subfamily (Figure 1). Detailed (sero)prevalences can be found in Supplementary Table 3. The most represented host genera were *Apodemus* (13 host species), *Oligoryzomys* (9 species), and *Akodon, Microtus*, and *Peromyscus* (6 species each). Host diversity was unevenly distributed across taxa, with *Apodemus agrarius* exhibiting the greatest orthohantavirus richness (4 virus species), while *Apodemus flavicollis, Apodemus sylvaticus, Akodon simulator, Microtus arvalis, Myodes glareolus, Oligoryzomys flavescens*, and *Sigmodon hispidus* each harbored 3 virus species. At broader taxonomic levels, *Apodemus* and *Microtus* were the most virus-rich genera (4 virus species each), and *Cricetidae* supported the highest overall virus diversity, containing 11 of the 12 orthohantavirus species represented in the dataset. Orthohantaviruses varied substantially in host breadth, with *Orthohantavirus hantanense* (commonly called Hantan virus) exhibiting the broadest host range (26 host species), followed by *Orthohantavirus andesense* (14 species; Andes virus), *Orthohantavirus tulaense* (9 species; Tula virus), and *Orthohantavirus dobravaense* (8 species; Belgrade-Dobrava virus). On the other hand, *Orthohantavirus chocloense* (Choclo virus) and *Orthohantavirus nigrorivense* (Nigrori virus) both only had one record each.

**Figure 1.**
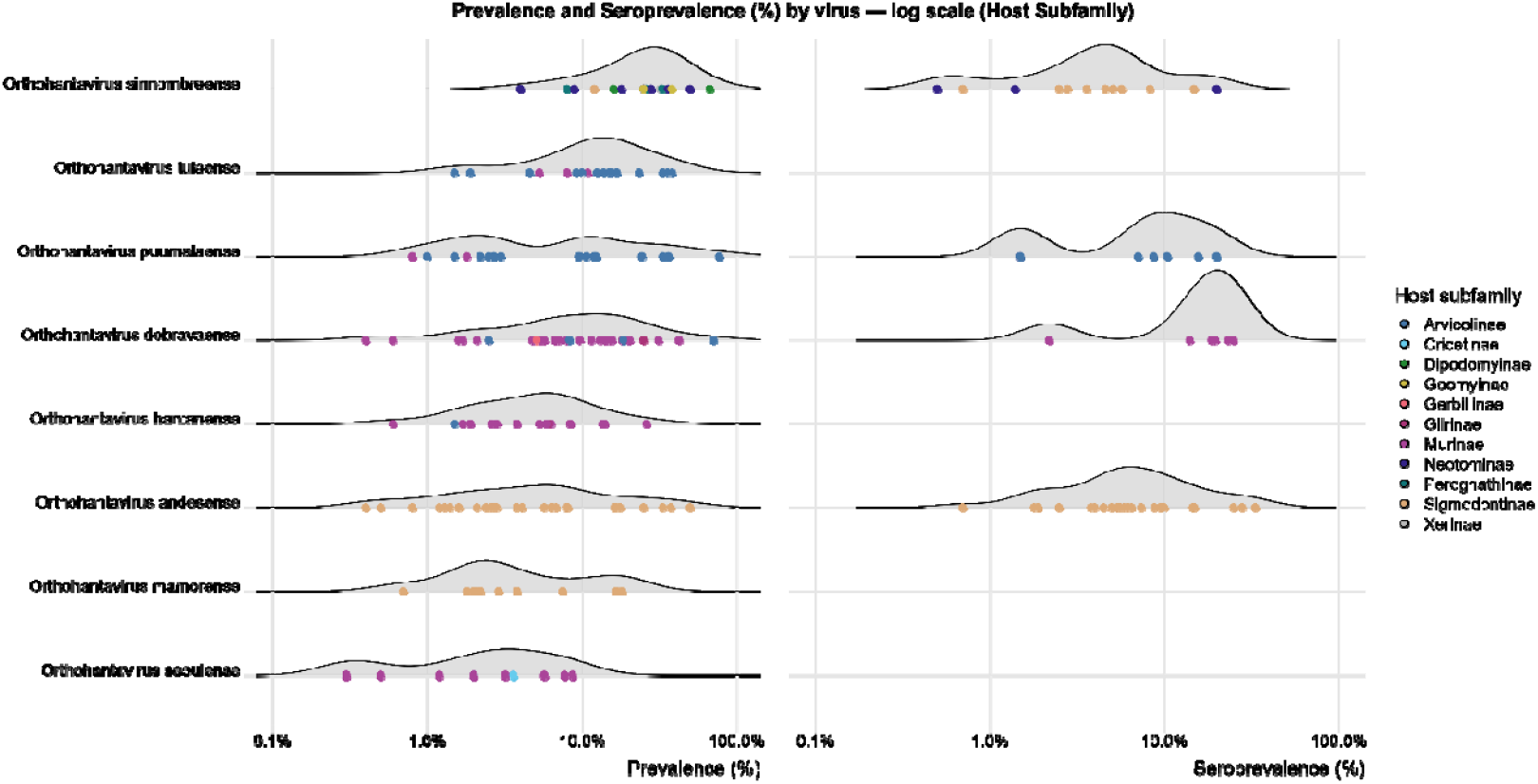
Global empirical distributions of Orthohantavirus prevalence and seroprevalence across key vertebrate reservoir subfamilies. Prevalences and seroprevalences are only shown for orthohantaviruses with more than 5 observations.

Studies included in the analysis sampled in North America, South America, Europe, and Asia, with only one source containing records from Africa and none from Australia. *O. andesense* and *O. mamorense*, had most of their records in South America (e.g. Argentina, Bolivia, Brazil, Chile, French Guiana, Uruguay, Paraguay, Peru). *O. bayoui, O. chocloense*, and O. *nigorivense* observations were based in North America (e.g. United States, Panama). *Orthohantavirus sinnombreense* (Sin Nombre virus) is most prevalent in both North and South America (Argentina, Brazil, United States). *O. dobravaense, Orthohantavirus puumalaense* (Puumala virus), and *O. tulaense* had observations reported mainly in Europe (Albania, Austria, Belgium, Bosnia, Bulgaria, Croatia, Czech Republic, Estonia, France, Germany, Greece, Hungary, Italy, Lithuania, Luxembourg, Montenegro, Netherlands, Poland, Romania, Russia, Serbia, Slovenia, Turkey, and Ukraine). Though, *O. puumalaense* also had a few observations from Asia. *O. hantanense, O*. seoulense, and *Orthohantavirus* thailandense (Thailand virus) were mainly found in Asia (China, Cambodia, Laos, Japan, Myanmar, North Korea, South Korea, Thailand, Vietnam) and Russia.

Detection of hantaviruses within reservoir populations exhibited heterogeneity within surveillance studies, with active viral prevalence (via PCR or viral isolation) and antibodies (seroprevalence) varying widely across host taxa and virus species (Figure 1). Prevalence estimates spanned nearly three orders of magnitude, ranging from a minimum of 0.3% (observed for *O. seoulense* in *Rattus tanezumi*) to an absolute maximum of 77.2% (*O. puumalaense* in *Myodes glareolus*) in some studies. The overall distribution of active prevalence was heavily right-skewed, with majority of host-virus associations clustered under 15%, while exceptional high-prevalence (>50%) were detected in specific hosts, such as *Microtus arvalis* harboring *O. dobravaense* (70.8%) and *Dipodomys spectabilis* harboring *O. sinnombreense* (67.0%). Seroprevalence data showed a similarly wide, yet distinct distribution, ranging from 0.5% (e.g., *O. tulaense* in *Apodemus sylvaticus*) to a maximum of 38.7% (*O. tulaense* in *Microtus subterraneus*). When disaggregated by host taxonomy, subfamilies displayed distinct epidemiologic profiles. For instance, members of the Arvicolinae and Sigmodontinae (family Cricetidae) frequently exhibited broader ranges and higher median active prevalence rates compared to the Murinae (family Muridae).

The systematically mapping these empirical distributions provides shows that orthohantaviruses are associated with a limited number of reservoir hosts, several lineages display broader host utilization across multiple rodent taxa, suggesting greater potential for host switching and geographic expansion and provides an ecological baseline to characterize the heterogeneity in host competence across rodent communities. Species consistently demonstrating elevated baseline prevalence can be distinguished from those exhibiting sporadic or marginal positivity, which may reflect transient spillover or localized, non-sustaining transmission pathways. Characterizing these distinct epidemiologic profiles ultimately refines species-specific hazard assessments, allowing for more targeted field surveillance and improved calibration of zoonotic spillover hazard models.

#### Climate driven shifts in global hantavirus spillover hazard

We trained and constructed climate driven model to predict the current and future spillover hazards through estimation of force of infection (FOI) through interactions of known rodent reservoirs of hantaviruses and human. Simulations predicted current hantavirus FOI, and future hantavirus FOI distribution for SSP 2-4.5 (moderate) and SSP 5-8.5 (severe) for years 2041-2060. Under the moderate climate change scenario (SSP 2-4.5, 2041-2060), the ensemble FOI models predicted a spatially uneven but geographically extensive increase in orthohantavirus spillover hazard relative to current conditions (Fig. 2, top left). Globally, mean ΔFOI across all grid cells was 0.095 (median = 0.059, SD = 0.160), with 74.9% of the modeled land area showing an increase in FOI (ΔFOI > 0), compared with 17.4% showing a decrease and 7.7% remaining effectively unchanged. The distribution of change was right-skewed, with the upper 5% of cells reaching ΔFOI ≥ 0.323 (95^th^ percentile), indicating that a subset of locations corresponding to the high-hazard belts identified in North America, Europe, and South America (Fig. 2, top left) are projected to experience disproportionately large increases in spillover hazard relative to the global average. Notably, the severe climate change scenario (SSP 5–8.5) produced an almost identical global signature (mean ΔFOI = 0.095; 74.9% of land area increasing; 95th percentile = 0.324), suggesting that by 2041-2060 the two emissions pathways have not yet meaningfully diverged in their net effect on global reservoir suitability, and that the difference between moderate and severe warming trajectories may only become apparent later in the century or in the spatial pattern rather than the magnitude of change. Models predicted that the modes pronounced increase orthohantavirus increase will be observed (Elevated FOI, Figure 2A) in three broad belts: (i) western and central North America, most pronounced across California and the Great Basin; (ii) western, central, and northeastern Europe, extending continuously into western Russia; and (iii) discrete patches across central South America, particularly the Amazon basin and southern Brazil. In contrast, areas of declining hazard (ΔFOI< 0) were spatially restricted, appearing mainly as isolated pockets bordering the high-ΔFOI belt in northeastern Europe and along limited stretches of the western US coast, suggesting these represent local contractions in reservoir suitability rather than a broad reorganization of hazard. The majority of the remaining global land area, most of Africa, South Asia, and Australia showed comparatively smaller change, indicating that projected hazard intensification under SSP 2-4.5 is concentrated in already-temperate reservoir ranges rather than emerging uniformly worldwide.

**Figure 2.**
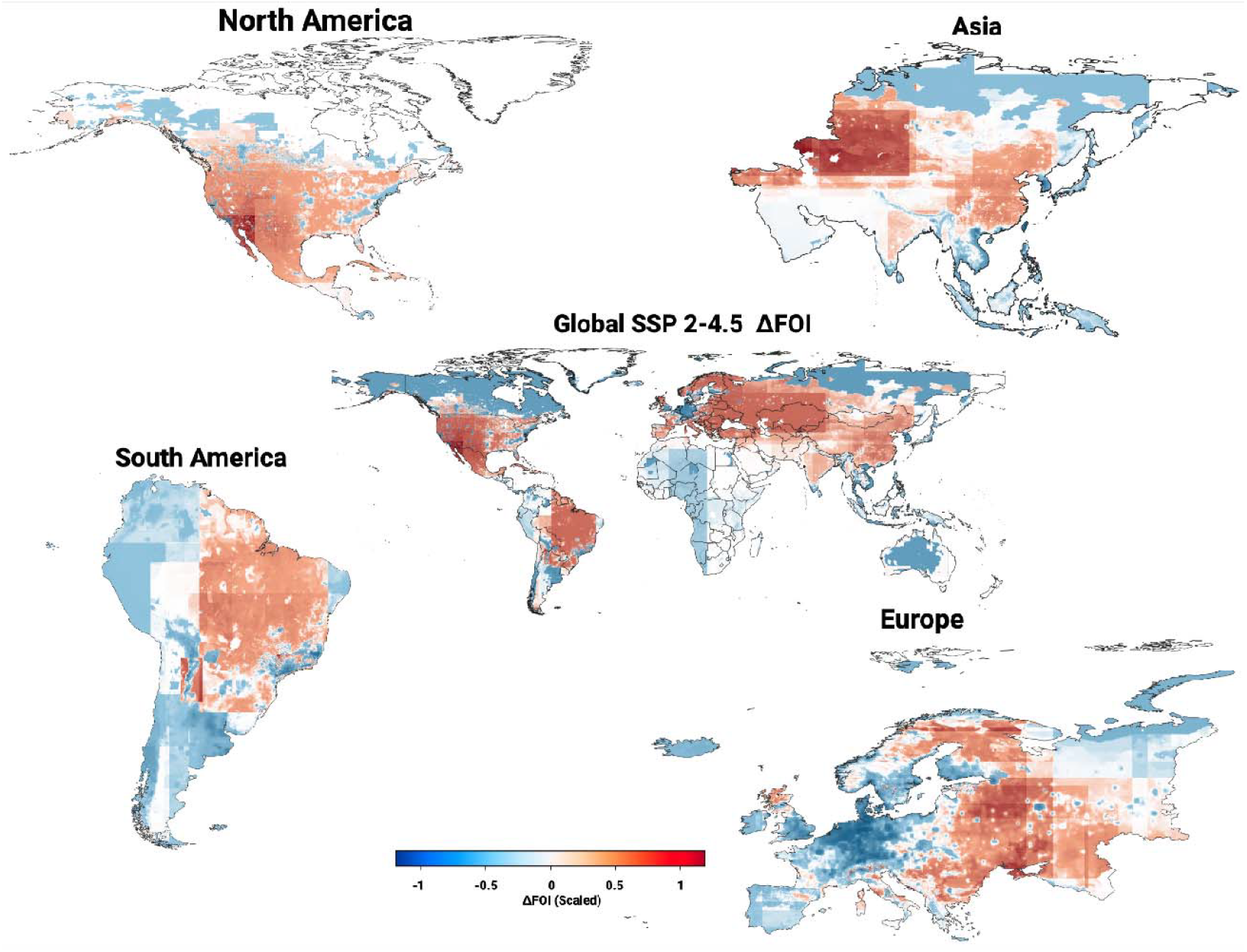
Climate-driven changes in orthohantavirus spillover hazard. Differences in force-of-infection (FOI) estimates between current conditions and future projections under SSP2–4.5 illustrate how climate-mediated shifts in reservoir suitability reshape zoonotic risk. Note: Scaling of Difference in FOI performed separately for global maps and each of the continents.

At the virus level, *O. sinnombreense* showed the largest projected mean increase in FOI under SSP2-4.5 (mean ΔFOI = 0.249), with 77.9% of its modeled range showing an increase, followed closely by *O. dobravaense* (mean ΔFOI = 0.243; 92.2% of its range increasing) and *O. tulaense* (mean ΔFOI = 0.240; 79.7% increasing). Notably, *O. dobravaense* combined a comparatively narrow range of change (min = – 1.468, max = 1.275) with the highest proportion of its range showing an increase of any virus modeled, indicating a spatially consistent, near-uniform intensification of hazard across its reservoir range rather than a pattern driven by a few extreme grid cells. By contrast, *O. sinnombreense* showed the widest spread of any virus (min = -3.128, max = 3.471), suggesting its elevated mean reflects a mix of strongly diverging local trajectories sharp increases in some areas offset by comparably sharp decreases in others consistent with the pronounced western North American hotspot identified in the global ΔFOI map (Fig. 2, top left). Viruses such as, *O. puumalaense, O. nigrorivense, O. bayoui*, and *O. hantanense*, showed moderate mean increases (ΔFOI = 0.08–0.11) despite high proportions of their ranges showing some positive change (65– 90%), indicating widespread but comparatively modest intensification. In contrast, *O. andesense, O. seoulense, O. mamorense*, and *O. thailandense* showed only marginal projected increases in mean FOI (ΔFOI = 0.03–0.05), and *O. chocloense* showed essentially no net change (mean ΔFOI = 0.0003), despite 71.0% of its range nominally showing an increase indicating that where change did occur, it was small in magnitude and largely offset by comparable decreases elsewhere. Virus-specific ΔFOI maps for all orthohantaviruses are provided in the Supplementary Material (Fig. S6 and S7).

Along with the increased spillover hazards, rich ness of zoonotic orthohantaviruses per grid under projected moderate climate change scenario (SSP2-4.5) showed the highest richness of 6–8 co-occurring hantaviruses concentrated across central and eastern Europe extending into western Russia, coincident with the region of highest ΔFOI. Along the Caribbean basin, Central America, and the Amazon and southern Brazil 4-6 orthohantaviruses were predicted. Bivariate maps in illustrate the regional distribution of orthohantaviruses FOI and projected virus richness in Americans (Figure 2C), Europe (Figure 2D) and Southeast Asia (Figure 2E). In the Americas, the bivariate map revealed a marked latitudinal divergence in hazards’ composition. Western and central North America, particularly California and the Great Basin/Intermountain West combined rising FOI with moderate-to-high richness, a pattern consistent with expanding habitat suitability for *Peromyscus* and *Sigmodon* reservoirs of Sin Nombre virus and related *O*.*sinnombreense-complex* viruses. Central America appeared as a transitional zone of moderate hazard and richness (yellow), while the Amazon basin and much of temperate South America fell in the low- ΔFOI/low-to-moderate richness quadrant (teal/green), indicating that the region’s projected hazard increase (Fig. 2, top left) is occurring against a comparatively sparse background of co-occurring hantaviruses largely reflecting the concentration of well-characterized Andes *(O*.*andesense)* and Mamoré virus *(Orthohantavirus mamorense)* reservoir associations rather than a multi-virus assemblage.

Europe showed the most spatially extensive and internally consistent high-risk signature of the three regions. A broad continuous band spanning the United Kingdom, Scandinavia, and eastern Europe into western Russia fell within the high-ΔFOI/high-richness quadrant, reflecting the combined expansion of climatically suitable habitat for Arvicolinae (notably *Myodes* and *Microtus*) and Murinae (*Apodemus*) reservoir taxa that carry Puumala and Dobrava-Belgrade viruses. Central Europe occupied an intermediate position (green/yellow), while Mediterranean margins trended toward lower joint risk, suggesting the intensifying hazard is not uniform across the continent but is instead weighted toward its northern and continental interior. In Southeast Asia, the bivariate signal was more spatially discrete. Southern China and adjacent mainland areas showed a strong high-ΔFOI/high-richness signature, consistent with the broad host range of *O. hantanense* and its principal reservoir, *Apodemus agrarius*. Coastal mainland Southeast Asia (Vietnam, Thailand, Myanmar) showed a moderate positive signal (pink), while island Southeast Asia (Indonesia, the Philippines) fell predominantly in the opposing, low-risk quadrant (blue), pointing to a sharper mainland–island contrast in projected hazard than is apparent in the Americas or Europe.

#### Environmental Drivers of Changing Spillover Hazard

To identify the underlying climatic variables governing projected shifts in spillover hazard we modeled associations between changes in FOI (ΔFOI) and changes in climatic and land-use variables in ensemble tree-based regression models. Permutation-based feature importance for the top five predictors of separately for each of the 11 Orthohantaviruses under both the SSP2-4.5 and SSP5-8.5 emission pathways (Figure 3 A) were computed.

**Figure 3.**
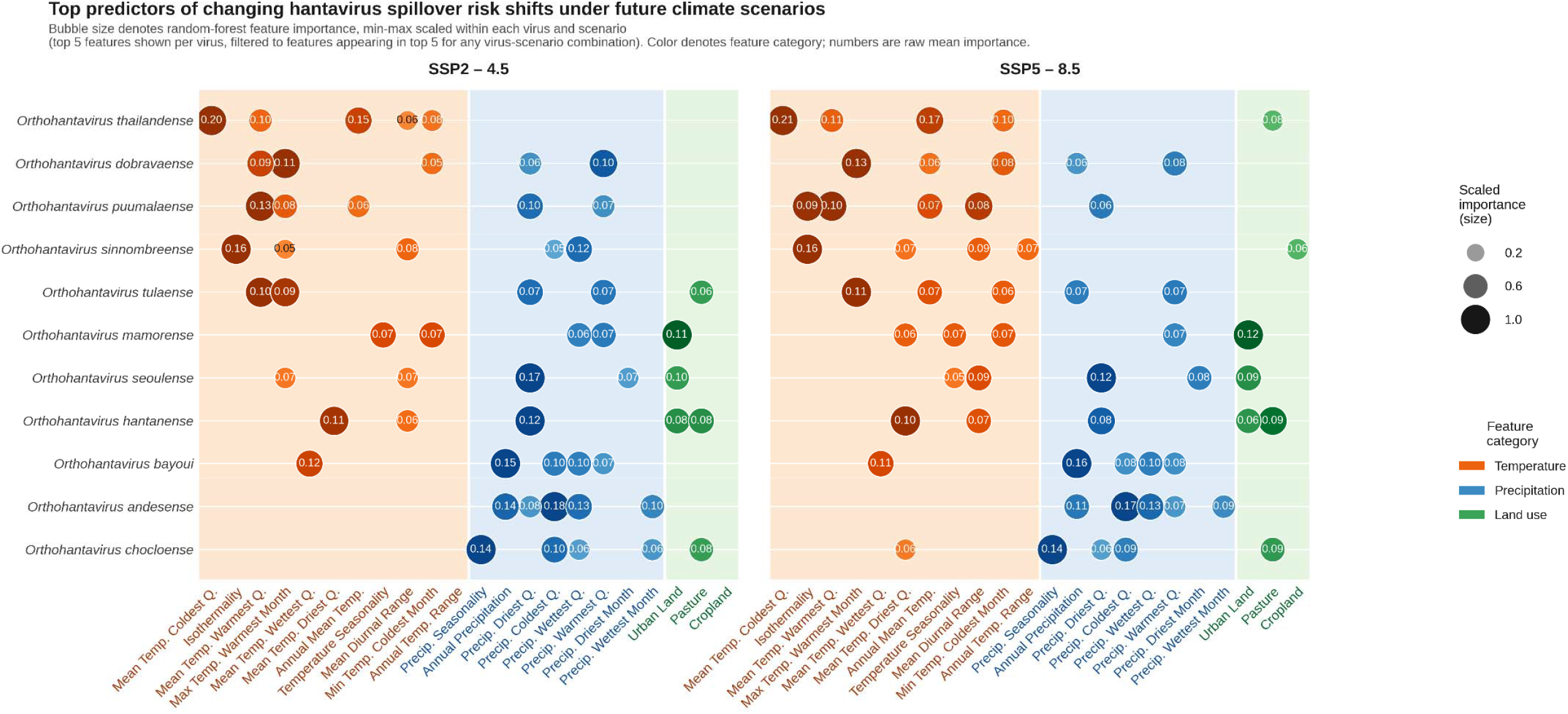
Top predictors of changing hantavirus spillover hazard shifts under future climate scenarios, Bubble size denotes random-forest feature importance, min-max scaled within each virus and scenario (top 5 features shown per virus). Color denotes feature category; numbers are raw mean importance.

##### Thermal niche constraints on future Orthohantavirus spillover hazards

Spillover changes for the first group of viruses demonstrated strong temperature sensitivity. *O. thailandense* emerged as a particularly striking temperature dependent outlier. In addition to *O. thailandense*, spillover hazard for *O. sinnombreense*, and *O. dobravaense* majorly showed high dependence on temperature related variables (Figure 3).

Top five important features for *O. thailandense* were all thermal related, with annual mean temperature as its leading predictor (mean scaled importance = 0.21 under SSP2–4.5 and 0.17 under SSP5–8.5, Figure 3 A). Partial dependence analyses revealed nonlinear, threshold-dependent responses to both mean temperature of the coldest quarter () and annual mean temperature (). In both cases, Force of Infection remained relatively stable below 2.2°C of warming, then increased steeply across a 0.4°C transition zone (2.2–2.6°C), before plateauing at higher temperature increases. This threshold-type response suggests a critical temperature tipping point for *O. thailandense* spillover hazard (Fig S5.9). The exceptional temperature dominance in *O. thailandense* indicates that future shifts in spillover hazard for this pathogen will be almost exclusively determined by warming trajectories rather than altered precipitation regimes or habitat conversion.

Partial dependence analyses for *O. sinnombreense* revealed strikingly negative associations with isothermality (Δ*Bio*_*3*_), whereby increases in temperature stability indicating more uniform seasonal temperatures decreased Force of Infection substantially (from ∼0.035 to <0.005 across the SSP2-4.5 range, Fig S5.9). This inverse pattern persisted under SSP5-8.5 with even steeper declines, suggesting that *O. sinnombreense* spillover hazard is maximized under thermally erratic conditions with high seasonal temperature variability. Precipitation of the wettest quarter showed equally strong negative dependence: increasing seasonal precipitation reduced FOI from ∼0.050 to ∼0.010, indicating this virus thrives in relatively dry conditions. Most strikingly, temperature annual range (Δ*Bio*_*7*_) exhibited a non-monotonic, U-shaped response across both scenarios, with optimal FOI at intermediate thermal ranges (∼0.8–1.0°C change) and elevated hazard at both extremes of annual temperature fluctuation.

*O. dobravaense* demonstrated warming-driven threshold responses coupled with precipitation sensitivity (Figure 3). Maximum temperature of the warmest month (Δ*Bio*_*5*_) and mean temperature of the warmest quarter (Δ*Bio*_*10*_) both exhibited steep sigmoid-shaped increases in FOI, with critical transitions occurring around 2.8–3.2°C warming under SSP2-4.5 and 3.0–3.5°C under SSP5-8.5 (Fig S5.4). Notably, precipitation of the warmest quarter showed a strong positive threshold effect (transition ∼-20 to 0 mm), indicating that increased wet-season precipitation combined with warming creates synergistically elevated spillover conditions. These patterns suggest *O. dobravaense* populations occupy ecological margins where concurrent warming and increased seasonal precipitation could rapidly expand suitable reservoir habitat.

##### Hydrological sensitivity to future Orthohantaviruses spillover hazards

*O. andesense, O. chocloense* and *O. bayoui* showed sensitivity to precipitation-driven environmental change, though with considerably more modest importance values than our previous estimates, suggesting more distributed ecological constraints across multiple bioclimatic and land-use dimensions (Figure 3 A).

Results for *O. andesense* indicated that changes in precipitation of the coldest quarter (Δ*Bio*_*19*_) showed a particularly striking threshold response, with FOI remaining near baseline for reductions of up to ∼15 mm, then increasing steeply across a narrow ∼15 mm transition zone (approximately −15 to 0 mm change) before plateauing at higher precipitation increases. Similarly, annual precipitation (Δ*Bio*_*12*_) and precipitation of the wettest quarter (Δ*Bio*_*16*_) both exhibited threshold dynamics, with the steepest increases occurring in the −50 to 0 mm and −20 to −10 mm ranges, respectively. These nonlinear responses suggest that *O. andesense* spillover hazard is sensitive to critical precipitation thresholds, with drying beyond these thresholds potentially driving increased viral transmission (Fig S 5.1).

*O. chocloense* exhibited more complex, distributed ecological constraints, showing negative associations with precipitation seasonality and land-use change intensity, along with a non-monotonic (bell-shaped) response to precipitation changes, suggesting that moderate precipitation levels rather than temperature or precipitation magnitude per se optimize transmission conditions for this species (Fig S5.3).

*O. bayoui* exhibited sensitivity to precipitation reduction, with partial dependence analyses revealing inverse relationships with both annual precipitation (Δ*Bio*_*12*_) and precipitation of the coldest quarter (Δ*Bio*_*19*_). Across both emission pathways, FOI decreased substantially as precipitation increased, suggesting that drying conditions rather than warming drive increased spillover hazard for this species. The virus also showed reduced sensitivity to mean temperature of the wettest quarter, indicating a complex ecological niche where moisture availability, rather than absolute temperature, constrains transmission (Fig S5.2).

##### Anthropogenic and landscape drivers of future Orthohantavirus spillover hazards

Another distinct profile emerged across several major viruses where land-use change functioned as a substantial driver of projected spillover hazard shifts. *O. seoulense, O. hantanense* and *O. mamorense* exhibited substantial land-use sensitivity alongside precipitation or temperature effects, indicating that habitat modification and anthropogenic landscape transformation represent critical climate-independent constraints on future reservoir suitability.

*O. seoulense* presented a fundamentally different profile characterized by precipitation seasonality sensitivity and unexpected urban land-use dependence. Precipitation of the driest month exhibited a strong negative relationship with FOI decreasing precipitation in dry seasons strongly predicted increased spillover hazard suggesting this virus thrives under water-stressed conditions. Urban land-use changes (ΔLU_Urban_) showed a pronounced positive threshold response, with FOI increasing steeply across the range of urbanization intensities. Mean diurnal temperature range (Δ*Bio*_*2*_) showed a monotonic negative relationship, indicating that reduced day-night temperature variability consistent with urban heat island effects or more stable microclimates favors transmission (Fig 5.8). This constellation of drivers implies that *O. seoulense* spillover hazard is driven less by large-scale climate patterns than by fine-scale urban habitat modification, where human settlement, reduced temperature volatility, and specific moisture regimes create optimal transmission conditions.

**Figure 4.**
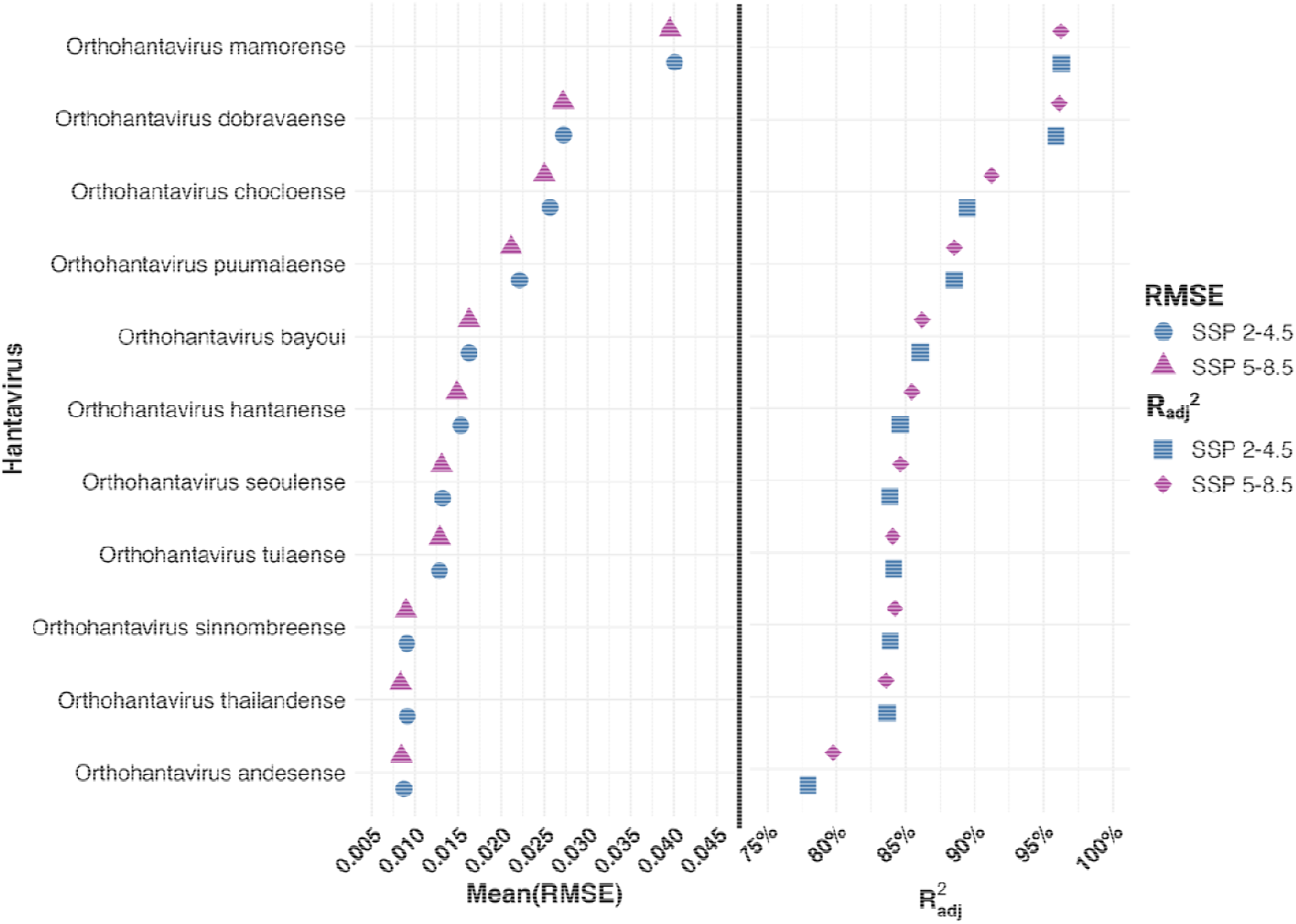
Performance metrics of models featuring average RMSE and Adjusted R^2^ values per Orthohantavirus in each of the two SSP scenarios.

**Fig 5.**
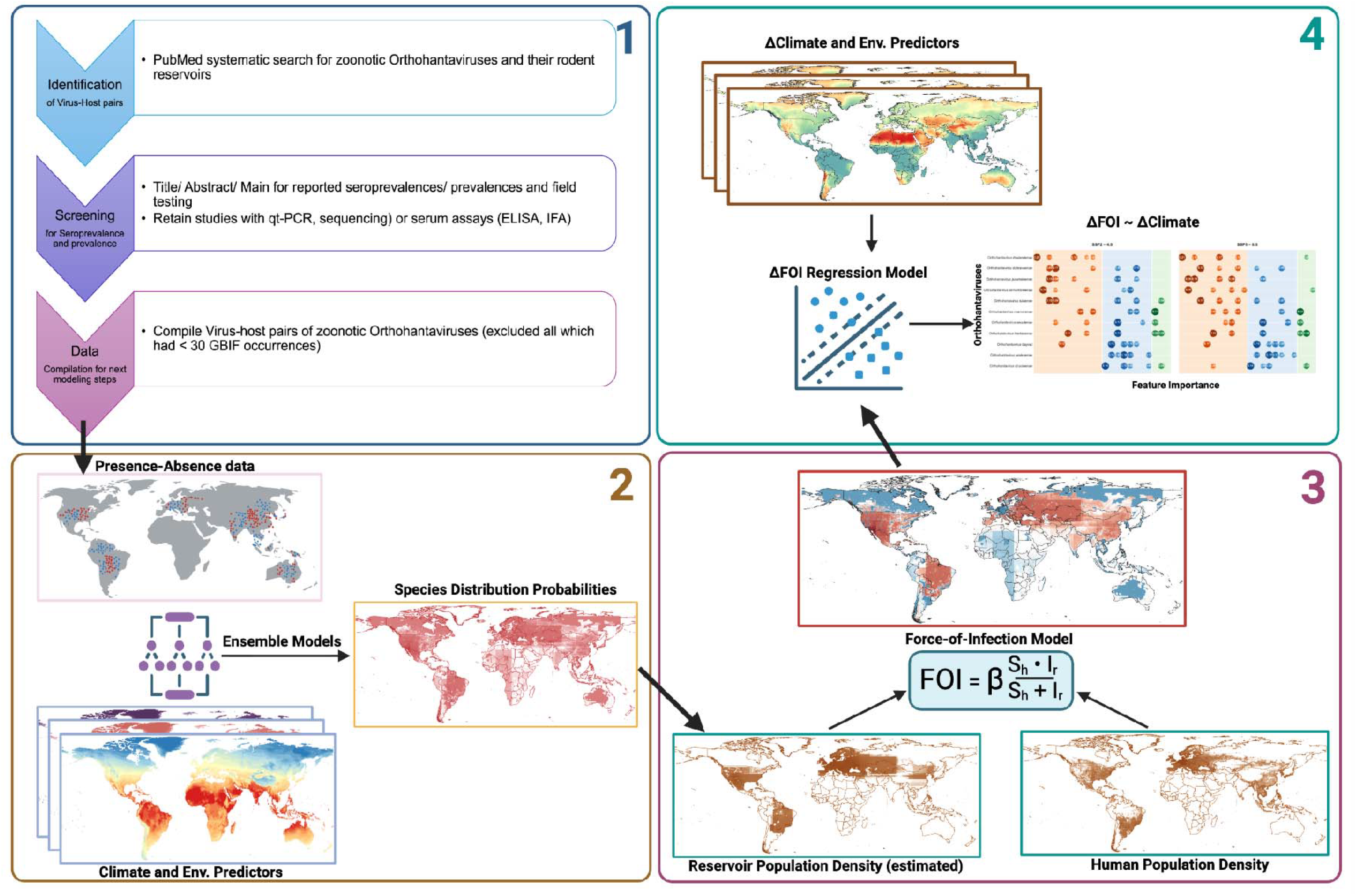
Methodological framework with (1) systematic literature search that informs (2) ensemble species distribution models, coupled with (3) a mechanistic force-of-infection formulation to (4) link changing climate and environmental suitability with disease risk in humans.

*O. hantanense* displayed the most complex, multi-dimensional response structure, combining temperature and precipitation seasonality with substantial land-use sensitivities. Mean temperature of the driest quarter (Δ*Bio*_*9*_) showed a strong positive threshold response (∼2–3°C warming range), while precipitation of the driest month exhibited the opposite: an inverse relationship where reduced dry-season precipitation increased FOI. Critically, land-use changes diverged sharply between scenarios: under SSP2-4.5, urban land use (Δ*LU*_*Urban*_) strongly increased FOI with a steep threshold response; under SSP5-8.5, pasture expansion (Δ*LU*_*Past*_) decreased FOI, suggesting that habitat composition and agricultural intensification may compete with optimal rodent reservoir conditions. These patterns indicate *O. hantanense* occupies an ecological niche where climate seasonality and anthropogenic landscape structure jointly determine spillover hazard, with the relative importance of urbanization versus agricultural land use varying contingently on the magnitude of climate change.

*O. mamorense* presented a distinctly urbanization-centric profile, with urban land-use change alongside temperature seasonality (exhibiting complex, non-monotonic U-shaped responses) emerging as comparably weighted drivers (Fig S5.6), suggesting that habitat modification and fine-scale thermal heterogeneity jointly determine spillover potential with minimal reliance on any single macroclimatic predictor.

##### Multidimensional ecological drivers of future Orthohantavirus spillover hazard

Two viruses *O. tulaense* and *O. puumalaense* showed mixed categories of feature importances (thermal, precipitation and landscape features) in top five important features predicting future spillover hazard (Figure 3).

*O. tulaense* showed distributed sensitivity across warming metrics and seasonal precipitation patterns, with comparably weighted dependencies on maximum temperature of the warmest month, mean temperature of the warmest quarter, and precipitation of the driest quarter that collectively intensify FOI across both climate scenarios, suggesting cumulative rather than threshold-driven climate effects (Fig S.5.11). *O. puumalaense* exhibited similarly dispersed feature importance across mean temperature of the warmest quarter, temperature seasonality, and precipitation dynamics, indicating an ecological generalist with FOI responding incrementally to multiple bioclimatic axes simultaneously rather than exhibiting single dominant environmental constraints.

Across the full virus panel, the relative macroecological influence of temperature, precipitation, and land-use on orthohantavirus spillover hazard appears to be scenario-independent, with comparable ranking structures under both SSP 2-4.5 and SSP 5-8.5 pathways. However, the prominence of land-use change as a driver alongside climatic variables suggests that managing spillover hazard in coming decades will require coordinated attention to both climate adaptation and spatial conservation planning that maintains habitat connectivity and minimizes anthropogenic disturbance in regions supporting high-hazard virus reservoirs.

The ΔFOI models were moderately robust across all the modeled viruses with a mean RMSE of 0.018 (0.008-0.04) and an average adjusted R^2^ value of 87% (78-96%) across both SSP scenarios with small standard errors (< 0.001) around the mean (Figure 3 B). The ΔFOI model *of O. thailandense* had the lowest mean RMSE (0.008, 0.009 for SSP 2-4.5, SSP 5-8.5 resp.) and the model of *O. sinnombreense* had the highest adjusted R^2^ score (96% each for SSP 2-4.5 and 5-8.5 resp.). Due to great variance in the total number of observations per model depending on the virus and the extent and the number of rodent reservoirs per virus(19K-1.88M), the adjusted R^2^ value differed substantially between each of the models.

## Discussion

Our analysis provides the first macroecological synthesis linking rodent reservoir ecology, prevalence heterogeneity and climate driven shifts in spillover hazards across full breadth of known zoonotic orthohantaviruses. The 2026 multi-country ANDV cluster illustrates how a long considered geographically confined virus can surface in and spread through limited human to human transmission chain events. Our models indicate that these events will occur frequently with increased initial spillovers at endemic niches of orthohantaviruses and their rodent reservoirs. Models for 12 zoonotic orthohantaviruses and their 61 reservoir species show that long term climatic changes in precipitation seasonality, increasing temperatures and precipitation will drive in increases predicted in spillover of risk of hantaviruses globally. These result differ significantly from ecology of arenaviruses and their reservoirs where land use changes were also shown to play an important role in changes in the force of infection^5^.

Projected changes in patterns associated with precipitation predominated signature for changing hazard of hantavirus spillover. These findings are broadly consistent with several decades of field ecology showing that rodent reservoir population dynamics and, in turn, hantavirus prevalence are governed less by mean climatic state than by the timing and magnitude of precipitation-driven resource pulses. The 1993 Sin Nombre virus outbreak in the southwestern United States, was preceded by an El Niño-associated precipitation anomaly that increased seed and vegetation productivity, driving a boom in deer mouse (*Peromyscus maniculatus*) density prior to the human outbreak^6^.

Our finding that precipitation of the driest and coldest quarters outweighs mean annual temperature as a predictor of ΔFOI for most New World viruses is consistent with this literature, and extends it by showing a similar, though mechanistically distinct, dry-season sensitivity among Old World lineages such as *O. seoulense* and *O. hantanense*. Field studies from Patagonia have shown comparable sensitivity of *Oligoryzomys longicaudatus*, the principal ANDV reservoir, to shifting precipitation trends, with modelled range contraction or eastward displacement under sustained drying^7^ a result that helps contextualize why

*O. andesense* in our models showed only a modest projected mean increase in FOI despite strong precipitation sensitivity: opposing local trajectories (range contraction in some areas, expansion in others) can offset one another in an aggregate metric, a pattern also evident in our results for *Orthohantavirus sinnombreense*.

The sharp, threshold-type warming responses identified for *O. thailandense* and *O. dobravaense* (Fig. 3) are consistent with the thermal physiology of their murine and arvicoline hosts. *Apodemus agrarius* populations show narrowing of the thermoneutral zone with local warming, such that individuals increasingly operate outside their zone of thermal comfort and must reallocate energy away from growth and reproduction once ambient temperature crosses population-specific critical points rodent body size often exhibits temperature-size patterns caused by differential thermoneutral zones and experienced extents of warming, with body size increasing and thermoneutral zone narrowing as temperature rises across space. This physiological threshold behavior offers a mechanistic explanation for the non-linear, tipping-point responses we observe in the partial dependence curves for thermally-sensitive viruses, rather than these being statistical artifacts of the ensemble regression^8^.

By contrast, the urban land-use dependence identified for *O. seoulense* and *O. mamorense* reflects a well-documented, largely climate-independent transmission pathway. *Rattus norvegicus* is the dominant reservoir of Seoul virus in dense urban settings globally, with seroprevalence and RNA-detection studies confirming sustained circulation in city rat populations from southern China^9^. *Rattus norvegicus* is the predominant reservoir of Seoul virus in urban residential areas in China, with population densities and community composition of rodents strongly influencing HFRS transmission to European urban parks a comprehensive serological and molecular characterization of Seoul virus in *Rattus norvegicus* in an urban park in Lyon, France confirmed continuous circulation with high seroprevalence and advocated for surveillance to be conducted at the scale of the entire city given the risk of transmission to humans^10^. That urbanization emerges as a top predictor independent of macroclimatic variables in our models reinforces the conclusion that Seoul virus risk will track patterns of informal settlement and rodent-proofing infrastructure as much as, or more than, climate trajectories.

The host-association data reveal a highly uneven landscape of knowledge. A small number of rodent genera *Apodemus, Microtus, Oligoryzomys* account for a disproportionate share of documented virus richness, while entire regions, most notably Africa, remain almost entirely unsampled despite growing molecular evidence of broader hantavirus circulation there. The projected shifts in force of infection are broadly consistent with the ecological logic established for other rodent-borne viral hemorrhagic fevers: increases in temperature seasonality and reductions in precipitation during key reservoir breeding periods emerged repeatedly among the top predictors of rising FOI, echoing the same climate variables identified as drivers of arenaviral and Lassa virus risk^11,12^. The concentration of high virus-overlap zones in temperate Europe and parts of Southeast Asia is particularly notable, as areas of high reservoir co-occurrence create conditions for multi-virus circulation within the same rodent communities and correspondingly more complex, harder-to-anticipate spillover dynamics. That several of the highest-magnitude projected increases (e.g., *O. thailandense, O. seoulense*) cluster in regions with dense human populations and agricultural land use reinforces the concern that climate change will not simply redistribute existing risk but may concentrate it at new human-reservoir interfaces.

The force-of-infection formulation relies on transmission rate parameter and reservoir density estimates. We present robust data to show estimation of these rate parameters. In rare occasions of absence of virus-specific empirical data, these parameters were informed by analogous rodent-borne systems, and it means absolute FOI values for those species-virus combinations should be interpreted as a relative risk surface rather than a calibrated probability of human infection. Reservoir probabilities occurrence from the ensemble SDMs further assume that current environmental relationships will hold under future climate and land-use regimes, a standard but non-trivial assumption for species with generalist habitat use. We addressed these uncertainties through repeated resampling, recursive feature elimination, and ensemble averaging across four tree-based algorithms, consistent with best-practice recommendations for propagating uncertainty in ensemble species distribution modeling; nonetheless, the resulting feature-importance rankings describe association, not causation identifying plausible ecological drivers.

Notably, in our literature review of prevalence and seroprevalence estimates for the FOI model, only one study covered a region in Africa (Table S3; Madagascar; Kikuchi et al., 2021)^13^. One reason for this is that many known African orthohantavirus reservoirs, including those from the Madagascar study, like *Mus musculus, Rattus norvegicus*, and *Rattus rattus*, were excluded from the literature review due to their ubiquitous global presences which would limit our model precision^14^. The ubiquitous presence of these hosts, alongside limited diagnostic capacity and likely underreporting of milder cases, exemplify key reasons for limited and inconsistent orthohantavirus prevalence data currently available from Africa^15^. Cross-sectional surveys of human orthohantavirus seroprevalence range widely from 0.2% to 17%^14^. Additionally, no records were found for Australia in the literature review, which is also consistent with previous studies. To date, Australia has had no reported human cases of orthohantavirus infections and very little evidence of orthohantavirus presence in Australian reservoirs^16,17^. More prevalence studies on orthohantavirus hosts need to be done in Africa and Australia in order to improve our model’s performance in these areas.

Where virus-specific transmission and reservoir density parameters were unavailable, we substituted biologically plausible estimates from analogous rodent-borne systems. This is a standard but non-trivial simplification in mechanistic spillover modeling: reviews of wildlife-to-human transmission modeling have repeatedly noted that zoonotic risk maps rely on multiple layers of statistical inference based on sparse data and simplifying assumptions, such that their main value lies in identifying blind spots in reservoir, pathogen, and human distribution data rather than in providing precisely calibrated hotspot predictions^18^. Similarly, for zoonotic infections with well-characterized spillover routes, the central modeling challenge is integrating statistical environmental suitability models with mechanistic models of population and infection dynamics precisely the coupling our framework attempts, and precisely where parameter substitution introduces the greatest uncertainty into absolute (rather than relative) FOI values The performance of our ΔFOI ensemble models (mean adjusted R^2^ = 87%, range 78–96%) compares favorably with other recent applications of ensemble machine-learning species distribution modeling to climate-driven zoonotic risk^19^. Similar tree-based ensemble frameworks combining random forest, boosted regression, and *MaxEnt* approaches have been used to project climate-driven shifts in habitat suitability for other zoonotic pathogens under SSP 2-4.5 and SSP 5-8.5, typically reporting AUC or TSS-based validation metrics in a comparable high-performance range an ensemble species distribution modelling approach integrating regression-based and machine-learning algorithms projected habitat suitability for *Coxiella burnetii* under SSP 2-4.5 and SSP 5-8.5 climate scenarios, evaluating performance using area under the curve and true skill statistics^20^. However, model performance metrics in this class of analysis primarily quantify goodness-of-fit to current environmental relationships rather than predictive skill under genuinely novel future climate space, an important caveat we return to below. The comparatively lower adjusted R^2^ for viruses such as *O. andesense* and *O. bayoui* is consistent with their smaller and more geographically restricted training datasets (19K–1.88M occurrence-informed grid cells across the panel), underscoring that model robustness in this framework is itself partly a function of underlying data density rather than ecological unpredictability.

Taken together, our findings indicate that climate change will not simply intensify hantavirus spillover hazard uniformly but will restructure it along virus-specific ecological axes thermal, hydrological, and anthropogenic that demand correspondingly differentiated surveillance and adaptation responses. This mirrors a broader pattern now documented across the mammalian virome: changes in climate and land use are projected to cause species to aggregate in new combinations at high elevations, biodiversity hotspots, and areas of high human population density in Asia and Africa, driving cross-species viral transmission an estimated 4,000 times by 2070, and this ecological transition may already be underway, with holding warming under 2°C unlikely to meaningfully reduce future viral sharing^21^. As the 2026 ANDV *(O. andesense)* cruise-ship cluster illustrates, geographically confined rodent-borne viruses can no longer be treated as static regional concerns; anticipating where and how orthohantavirus reservoir suitability will shift is a prerequisite, not a luxury, for global pandemic preparedness.

## Methods

We extended well established and validated framework for climate-driven zoonotic risk assessment developed by Kulkarni et al. to quantify global patterns of orthohantavirus spillover hazard^5,22^. This approach integrates ensemble machine learning–based species distribution models (SDMs) with a mechanistic force-of-infection (FOI) formulation to link environmental suitability of reservoir hosts with human exposure dynamics. SDMs were implemented using an ensemble of tree-based algorithms within the Python *mlsdm* framework^22^, enabling robust estimation of reservoir occurrence probabilities while accounting for nonlinear responses, multicollinearity, and complex interactions among climatic and land-use variables (Figure 5). Reservoir distributions were then coupled with probabilistic representations of infection prevalence within estimated rodent population densities and the human population density to estimate spatially explicit FOI as a proxy for spillover hazard. Model development incorporated repeated cross-validation, pseudo-absence generation, recursive feature selection, and ensemble averaging to ensure generalizability and minimize overfitting. Uncertainty was propagated through iterative resampling and stochastic parameterization, and associations between environmental change and spillover hazard were quantified using ensemble regression and permutation-based feature importance analyses. Together, this integrated framework provides a scalable and reproducible approach for predicting zoonotic risk under current and future climate scenarios.

### Compilation of orthohantavirus-host association data and reservoir determination

We compiled a comprehensive database of orthohantavirus–host associations through systematic aggregation of published literature, curated databases, and publicly available published datasets^23,24^. We started by focusing on 12 different orthohantaviruses associated with rodent-human transmission: *Orthohantavirus andesense, Orthohantavirus bayoui, Orthohantavirus chocloense, Orthohantavirus dobravaense, Orthohantavirus hantanense, Orthohantavirus mamorense, Orthohantavirus nigorivense, Orthohantavirus puumalaense, Orthohantavirus seoulense, Orthohantavirus sinnombreense, Orthohantavirus thailandense*, and *Orthohantavirus tulaense*. We initially focused on searching virus-host associations previous published by Pandit et al 2021^23^ and included new associations found through our searches.

This systematic review was conducted primarily through PubMed by using the search terms “(*any of the orthohantavirus names*) AND (prevalence) AND (*any of the scientific host names*)”. Alternative virus and host names are based off of names listed in the International Committee on Taxonomy of Viruses (ICTV) and the American Society of Mammalogists Mammal Diversity Database, respectively, along with results found during our systematic review. To supplement host-virus combinations with few results from PubMed, we would search Google Scholar using the terms ““(*any of the orthohantavirus names*)” prevalence “(*any of the scientific host names)*”“.

Records were included if they documented viral detection in non-human vertebrates using molecular (RT-PCR), virological (virus isolation), or serological (e.g., ELISA, immunofluorescence) assays. Host–virus associations were retained only when supported by primary evidence and accompanied by sufficient metadata, including geographic location. The full texts of articles were screened for inclusion, and information was then compiled on studies’ reported prevalence or seroprevalence data, sample sizes, study areas, and viral detection methods. For virus-host combinations with many results, newer publications and publications providing prevalence data (e.g. using RT-PCR, viral isolation) were prioritized. Publications with 100% prevalences or seroprevalences were excluded due to the likelihood of sampling bias. Details of the included publications and other metadata collected are presented in the Supplementary Table 3.

We adopted a hierarchical framework extending established approaches used in rodent-borne zoonoses to differentiate if rodent reservoirs are natural reservoirs of the orthohantavirus. Host species were classified as reservoirs if they met at least two of the following criteria: (i) repeated detection of viral RNA or antigen across independent studies or geographic locations; (ii) evidence of active infection (PCR-positive individuals), rather than solely seropositivity; (iii) ecological plausibility for sustained intraspecific transmission, including species abundance, habitat suitability, and known host specificity of orthohantaviruses; and (iv) documented association with human cases or outbreak settings that were spatially and temporally consistent with spillover events. When multiple species met these criteria for a given virus, all were retained to capture potential multi-host transmission systems. Prevalence and seroprevalence data were compiled where available and used to inform distributions of infection probability within reservoir populations (see modeling FOI section). For species lacking prevalence data, infection probabilities were estimated probabilistically using binomial approximations to account for heterogeneous sampling effort.

### Modelling datasets

Georeferenced historical occurrence records for confirmed and putative reservoir species were obtained primarily from the Global Biodiversity Information Facility (GBIF)^25^. Records were cleaned to remove duplicates and erroneous coordinates and were restricted to observations from 1990 onward to align with environmental covariates. Presence-only records were converted into presence–absence datasets using environmentally stratified pseudo-absence sampling to reduce sampling bias and improve representation of environmental space.

We assembled a suite of environmental predictors capturing climatic, ecological, and anthropogenic drivers of reservoir distributions. These included 19 bioclimatic variables^19^ (temperature and precipitation), elevation (DEM)^26^, and land-use variables^27^ such as cropland, urban, secondary land, forest, and grassland cover. All raster datasets were resampled to a common spatial resolution of 0.042° and aligned spatially (WGS84) to ensure comparability across datasets. Future projections of environmental variables were obtained from CMIP6^28^ (Coupled Model Intercomparison Project Phase 6) under Shared Socioeconomic Pathways: SSP 2-4.5 (moderate/ middle of the road climate change scenario) and SSP 5–8.5 (Extreme climate change scenario) for the period 2041–2060. Raster stacks were harmonized across current and future scenarios to ensure consistency.

### Species distribution modeling (SDM)

We estimated the probability of occurrence of each reservoir species using an ensemble species distribution modeling framework. To capture the complex and non-monotonic relationships between environmental drivers and species distributions, we employed tree-based machine learning algorithms, including random forests, extra trees, extreme gradient boosting, and light gradient boosting machines. Models were trained using repeated resampling with a 75:25% train–test split and fivefold cross-validation. To improve robustness and account for sampling variability, the modeling workflow was iterated across multiple pseudo-absence realizations (n = 100/rodent species), and predictions were averaged across algorithms and iterations. Recursive feature elimination with five-fold stratified cross-validation was used to select informative predictors and mitigate overfitting and multicollinearity among environmental variables. Model performance was evaluated using cross-validation and independent test-set metrics, namely, accuracy, precision, recall, AUC score. Ensemble averaging allowed propagation of uncertainty and to avoid reliance on single-model outputs. Model training was conducted by modifying the functions from *mlsdm* python package^22^.

### Estimation of zoonotic spillover hazard

We quantified zoonotic spillover hazard using a mechanistic proxy metric, the force of infection (FOI), representing the rate of effective contact between infectious reservoir hosts and susceptible humans. FOI was modeled as a density-dependent function incorporating predicted reservoir presence probabilities, estimated reservoir infection probabilities, and human population density. In the absence of direct data on reservoir densities and transmission parameters for all orthohantaviruses, model parameters were informed by biologically plausible estimates derived from analogous rodent-borne systems. Reservoir abundance within each grid cell was approximated probabilistically using SDM-derived occurrence probabilities (*p*_*i*_)and infection status was simulated using binomial sampling informed by observed or inferred seroprevalence ranges (*π*). FOI was calculated for each reservoir species and aggregated across all reservoirs for a given virus. Spatial FOI surfaces were generated for current conditions and future climate scenarios, and differences between scenarios were used to quantify projected changes in spillover hazard.

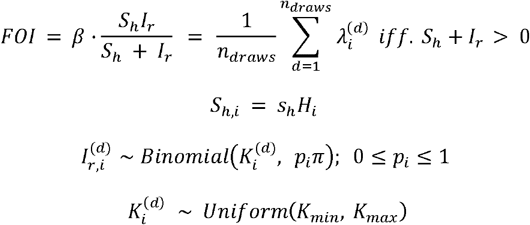

Where, (*S*_*h*_)was the product of human population density and proportion of susceptible individuals in the *S*_*h*_ population in each grid cell *i, I*_*r*_was the estimated infectious reservoir density based on reservoir presence 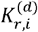 in the same grid cell *i* that integrated the seroprevalence *π*, expected range of rodents [*K*_*min*,_ *K*_*max*_]present in each grid cell and the probabilities of presence from the SDM *p*_*i*_. The force-of-infection 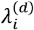 was calculated stochastically for (*d*) draws and averaged over all *n* _*draws*_ to give overall *FOI* for each grid cell per rodent species. 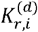 was drawn from estimated family size and rodent densities based on key ecological and life history traits using machine learning algorithms (for details, see Han et al. 2020^29^).

### Association of spillover hazards with environmental drivers

To examine the drivers of projected changes in spillover hazard, we modeled associations between changes in FOI and changes in climatic and land-use variables. Differences between current and future conditions were computed for both FOI (Δ *FOI*) and predictor variables (Δ *ssp*) and used as inputs in ensemble tree-based regression models. Feature importance was quantified using permutation-based metrics across repeated model fits, and partial dependence functions were used to characterize nonlinear relationships between key predictors and changes in FOI. This approach captured higher-order interactions and avoided assumptions of linearity. Of the 12 *Orthohantavirus* under study, 11 were retained for building these models (*O. nigorivense* was excluded due to lack of sufficient variation in data on Δ *FOI* and. Δ *ssp*) In total, 22 models (11 viruses X 2 SSP scenarios) were fit for capturing the association of spillover hazard with environmental drivers.

## Data Availability statement

All the data used in this manuscript comes from open-source databases and has been cited in appropriate places within the text of the manuscript. All the code and generated metadata to perform the analyses mentioned in this manuscript are available at https://github.com/EpiPandit/Pandit_et_al_orthohantavirus. Figures 2, 4 and 5 in this manuscript were generated in R v4.4.0 (https://www.R-project.org/) and arranged in BioRender (© 2026; https://BioRender.com).

## Acknowledgements and Funding

Development of the analytical and modeling frameworks used in this manuscript was supported by the Wellcome Trust (grant number 226099/Z/22/Z). P.S.P. acknowledges additional support from the National Science Foundation under Award Number DMS-2325267.

