## Supplementary Material for "Ecological Cascades and Future Hantavirus Spillover Risk in a Changing Climate"

Tables and Figures

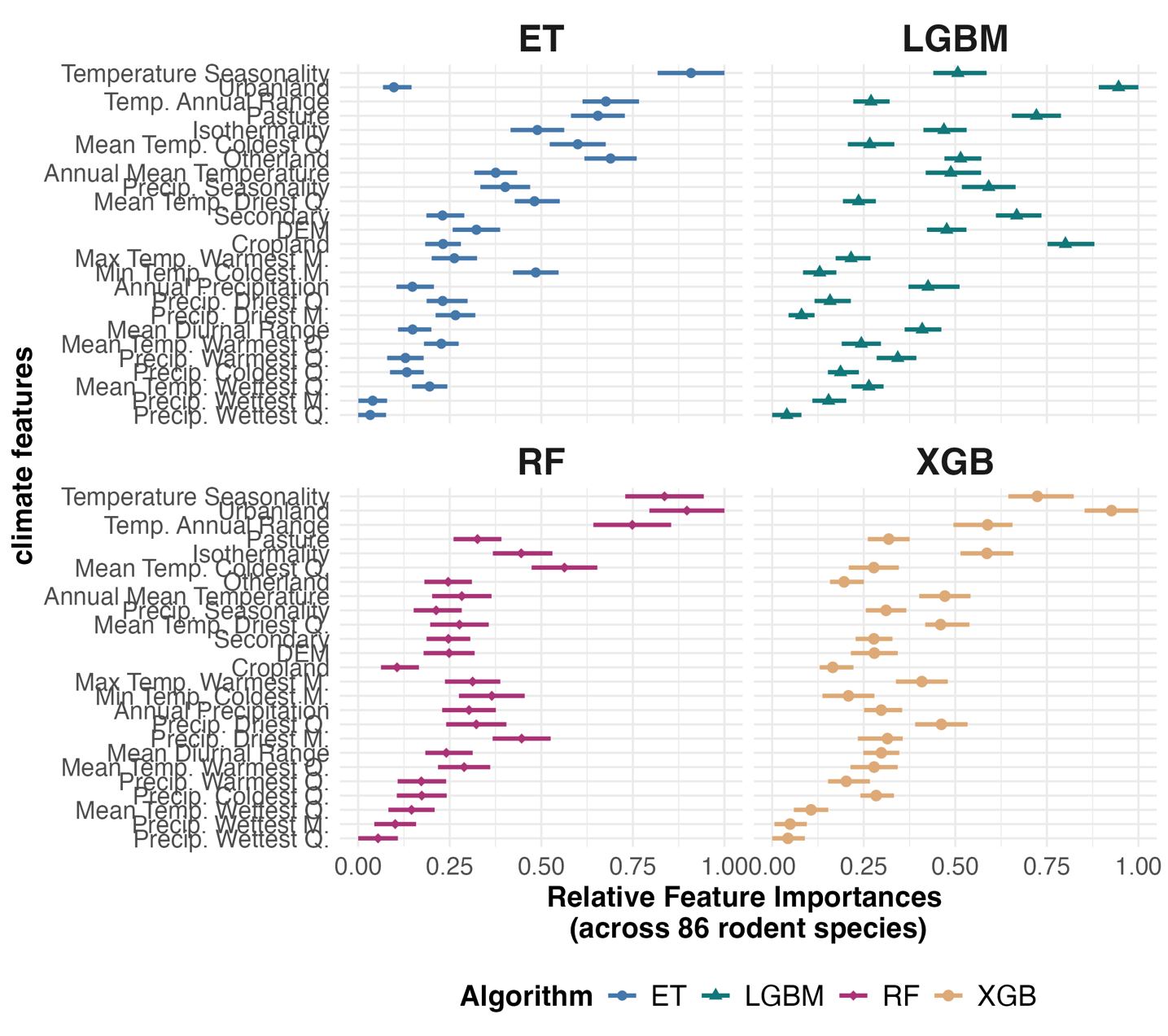

Fig S1. Relative importance of climate, environmental and land use features in predicting distribution probabilities of Orthohantavirus rodent reservoirs. Note: The figure illustrates the mean relative feature importance (between 0 and 1) for all 25 features across 61 rodent species.

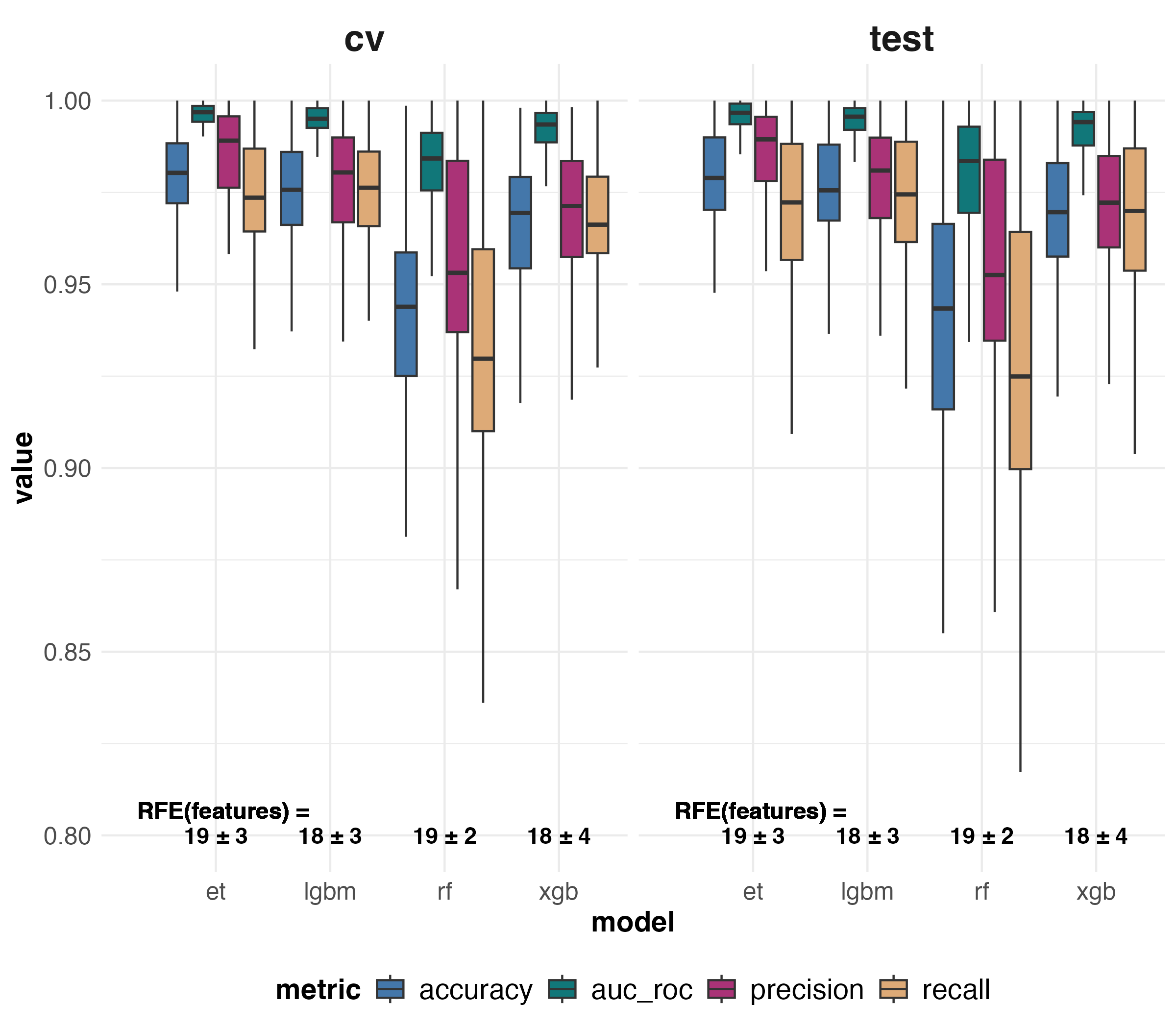

Fig S2. Summary of performance metrics from Species Distribution Modeling of 61 Orthohantavirus rodent reservoirs with Recursive Feature Elimination (RFE) applied over 100 iterations for each rodent species.

Table S1. Spatial correlation within and between the estimated Force-of-Infection (FOI) maps calculated with Pearson’s test of correlation and Moran’s I test (averaged for all 61 Orthohantavirus rodent reservoirs).

| **Pearson’s correlation (Mean ± SD)** | | **Moran’ I statistic (Mean ± SD)** | |
| --- | --- | --- | --- |
| Corr (Current, SSP 2-4.5) | 0.21 ± 0.15 | Current | 0.95 ± 0.06 |
| Corr (Current, SSP 5-8.5) | 0.20 ± 0.15 | SSP 2-4.5 | 0.79 ± 0.14 |
| Corr (SSP 2-4.5, 5-8.5) | 0.88 ± 0.15 | SSP 5-8.5 | 0.79 ± 0.14 |
|  |  | SSP 2 – Current | -0.16 ± 0.16 |
|  |  | SSP 5 – Current | -0.16 ± 0.16 |
|  |  | SSP 5 – SSP 2 | 0.002 ± 0.007 |

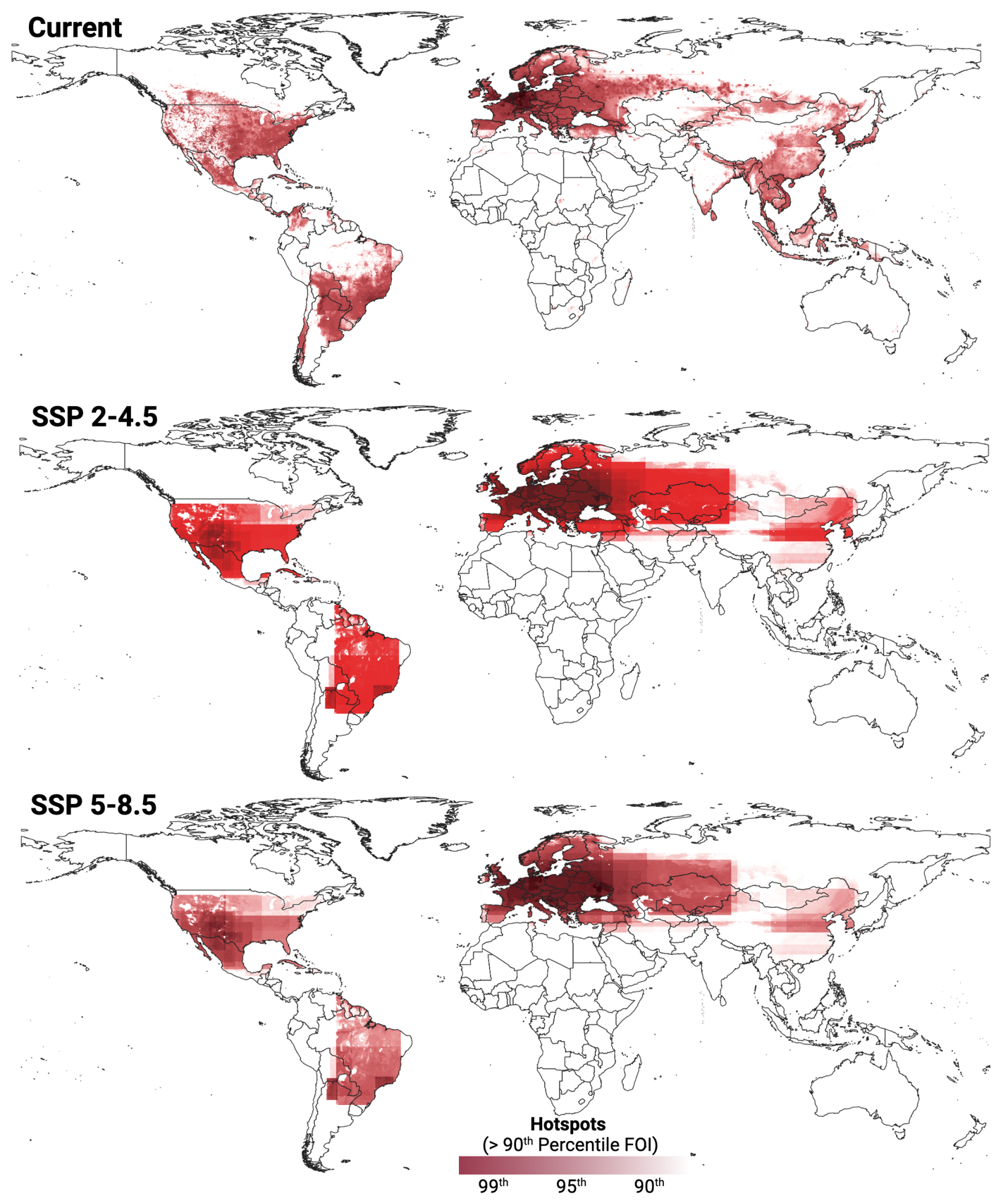

Fig S3. Potential hotspots for outbreak of Orthohantavirus based on estimated Force-of-Infection (FOI) metric. Areas were designated as hotspots where FOI was estimated in the 90-99^th^ percentile; averaged over all 12 modeled Orthohantaviruses.

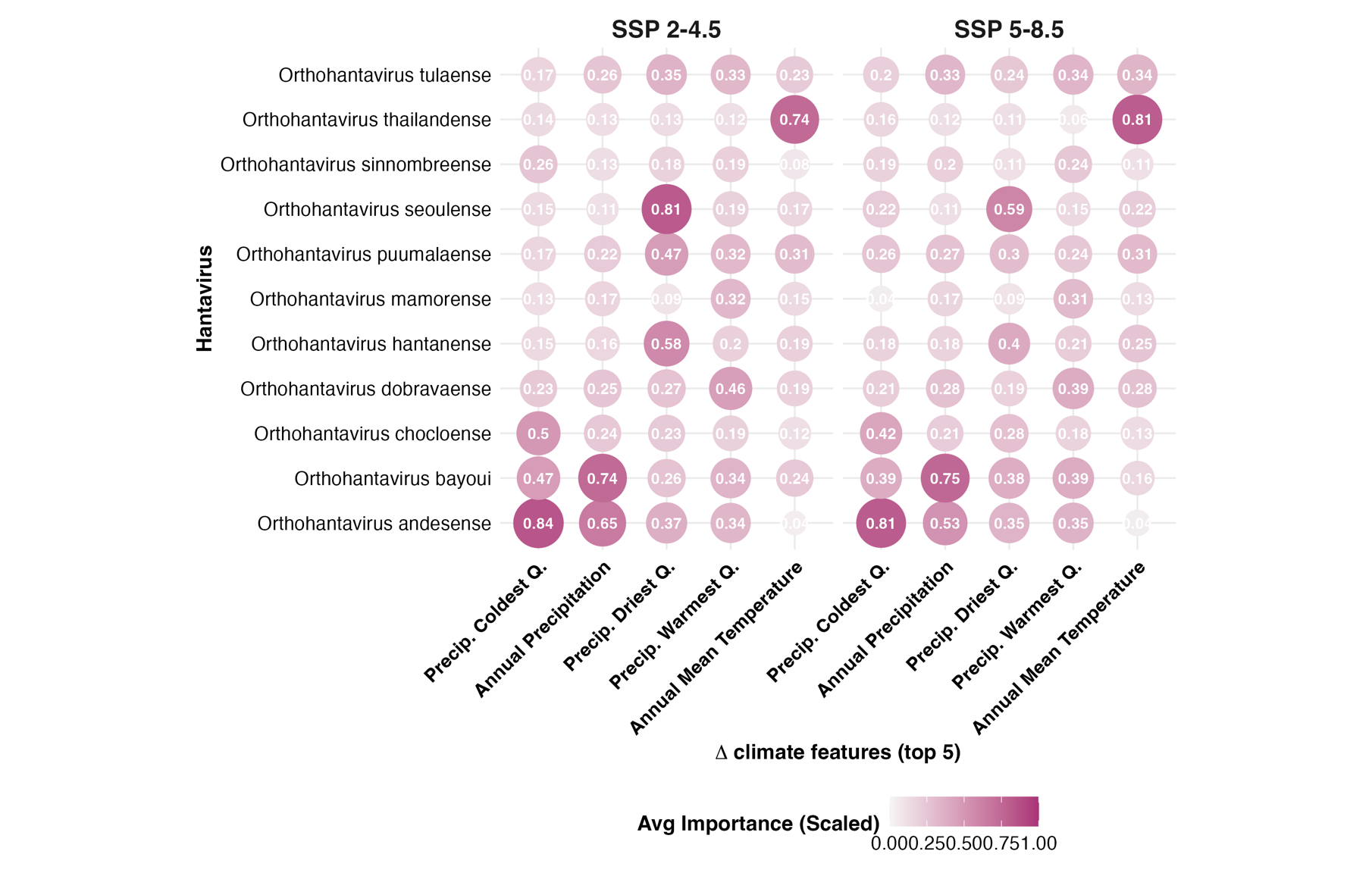

Fig S4. Feature importances of top 5 overall features across all 11 modeled Orthohantaviruses.

### Partial dependence plots of $\boldsymbol{\Delta}\boldsymbol{FOI}$ models for top three features per virus and SSP scenario combination

Table S2. Glossary of terms in the following plots

| **Name in graph** | **Description (Difference between SSPs and Current climate scenario of –)** |
| --- | --- |
| Delta_bio_1 | Annual Mean Temperature |
| Delta_bio_2 | Mean Diurnal Range |
| Delta_bio_3 | Isothermality |
| Delta_bio_4 | Temperature Seasonality |
| Delta_bio_5 | Max Temperature of Warmest Month |
| Delta_bio_6 | Min Temperature of Coldest Month |
| Delta_bio_7 | Temperature Annual Range |
| Delta_bio_8 | Mean Temperature of Wettest Quarter |
| Delta_bio_9 | Mean Temperature of Driest Quarter |
| Delta_bio_10 | Mean Temperature of Warmest Quarter |
| Delta_bio_11 | Mean Temperature of Coldest Quarter |
| Delta_bio_12 | Annual Precipitation |
| Delta_bio_13 | Precipitation of Wettest Month |
| Delta_bio_14 | Precipitation of Driest Month |
| Delta_bio_15 | Precipitation Seasonality |
| Delta_bio_16 | Precipitation of Wettest Quarter |
| Delta_bio_17 | Precipitation of Driest Quarter |
| Delta_bio_18 | Precipitation of Warmest Quarter |
| Delta_bio_19 | Precipitation of Coldest Quarter |
| Delta_lu_crop | Land use – Cropland |
| Delta_lu_urbn | Land use – Urban land |
| Delta_lu_pastr | Land use – Pasture |
| Delta_lu_secd | Land use – Secondary land |
| Delta_lu_othr | Land use – Other/ Barren land |
| Delta_DEM | Elevation/ Digital Elevation Model |

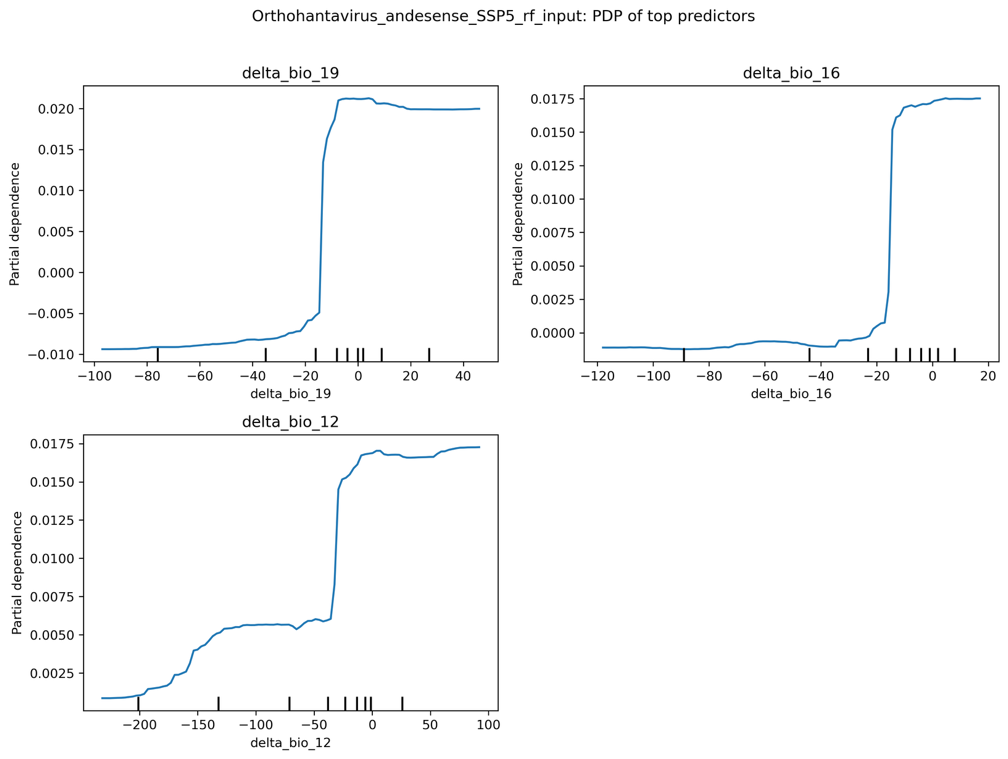

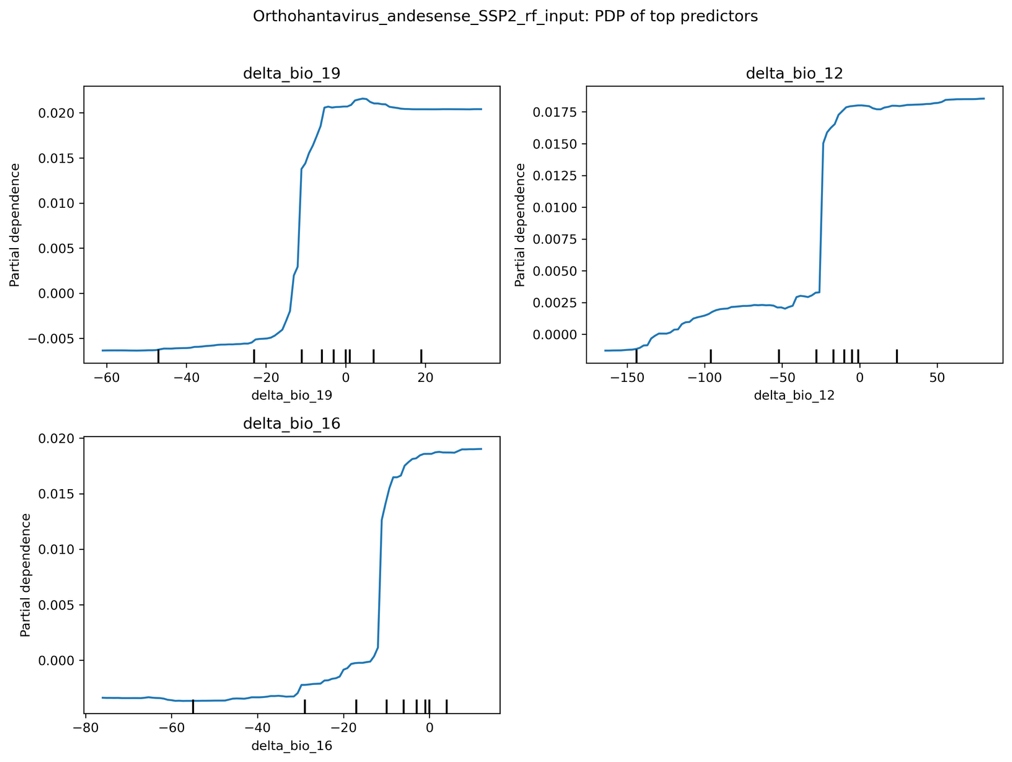

Fig S5.1. Orthohantavirus andesense

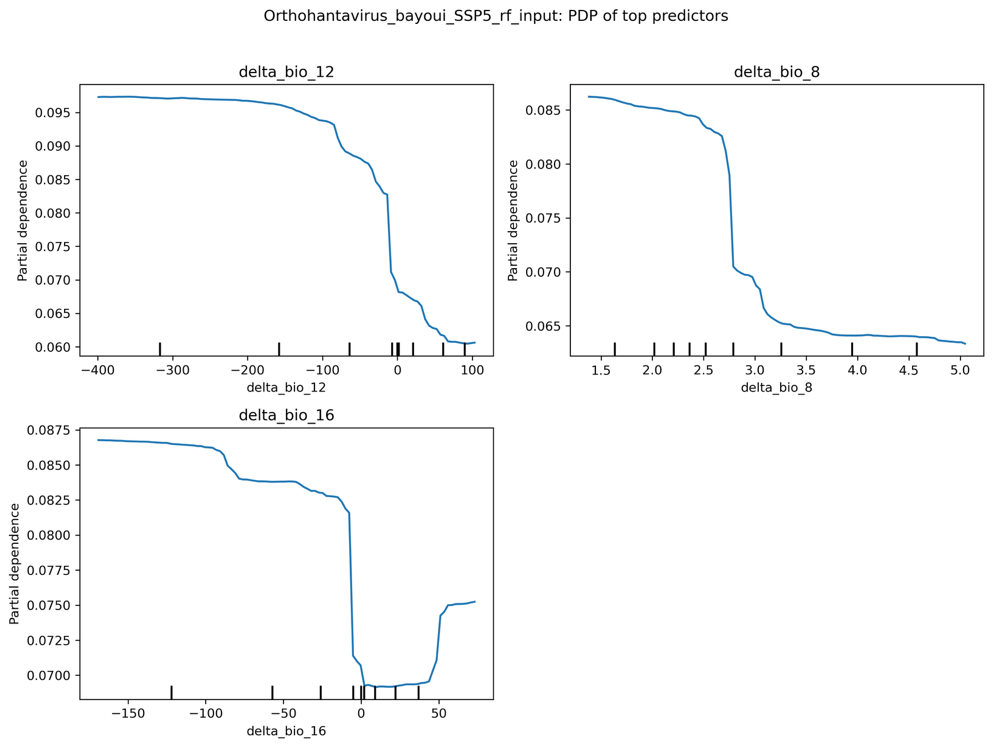

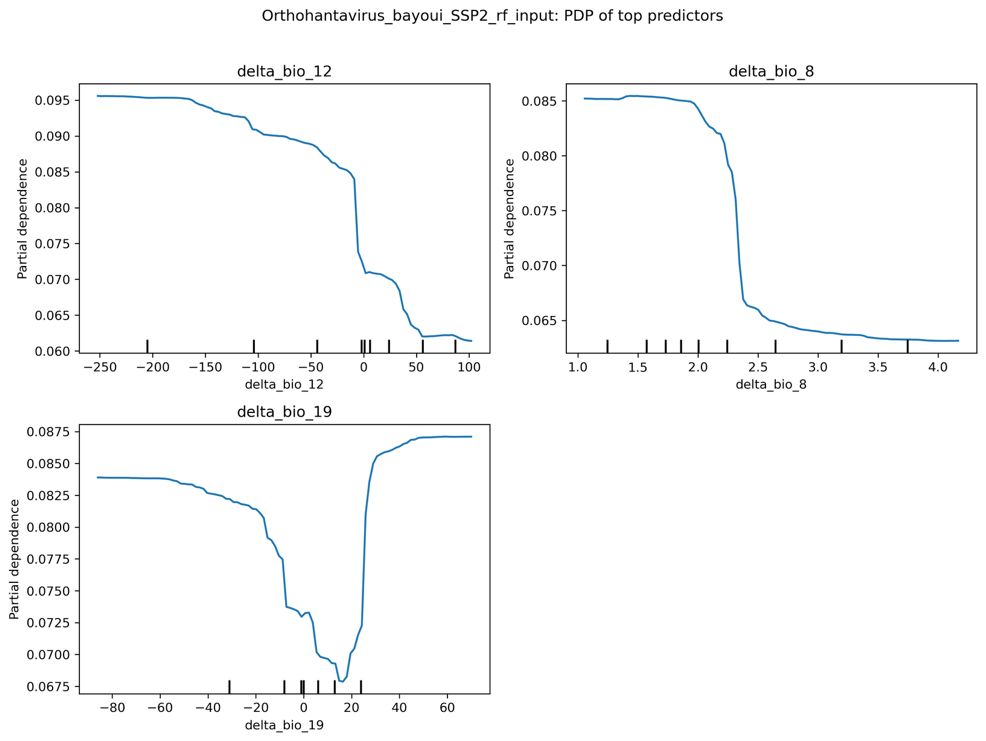

Fig S5.2 Orthohantavirus bayoui

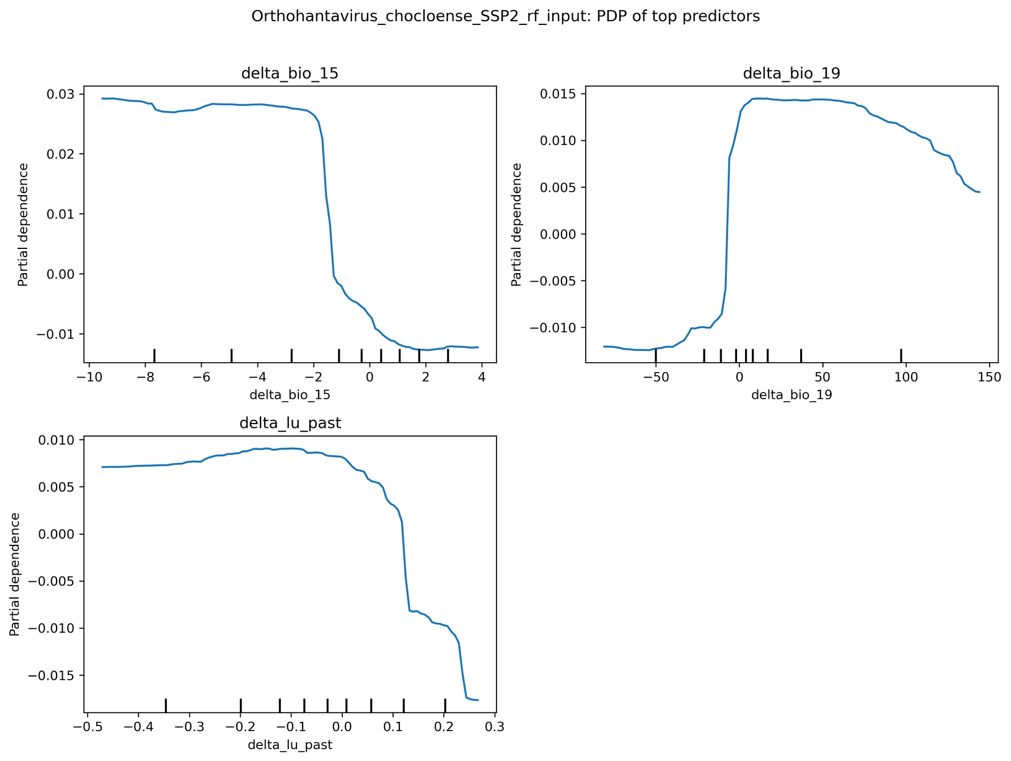

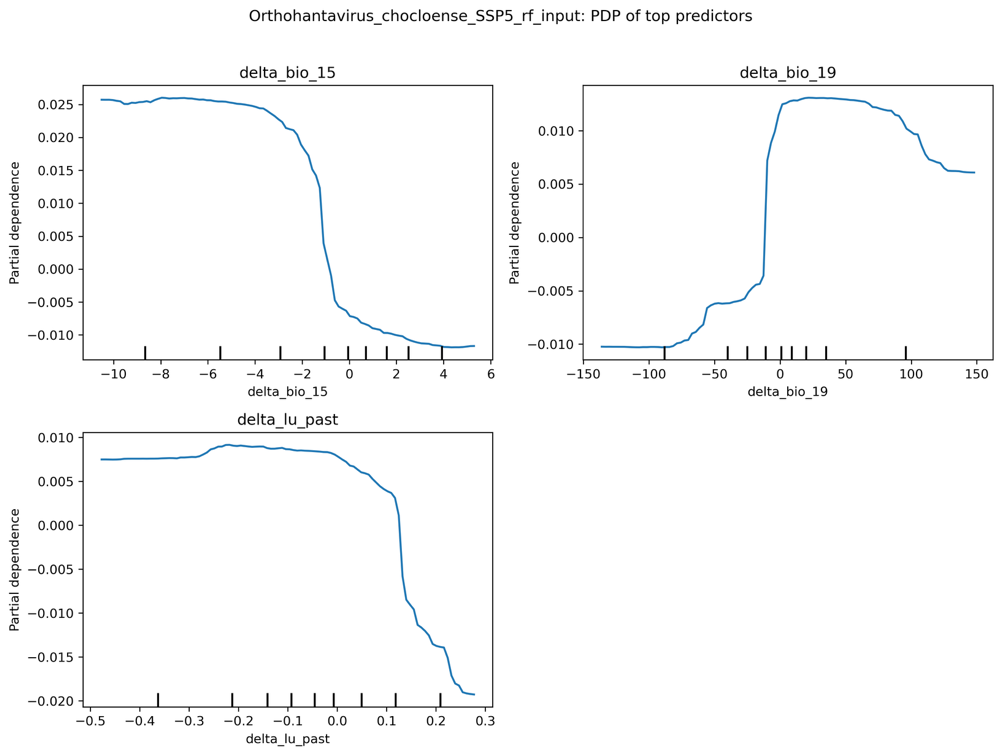

Fig S5.3. Orthohantavirus chocloense

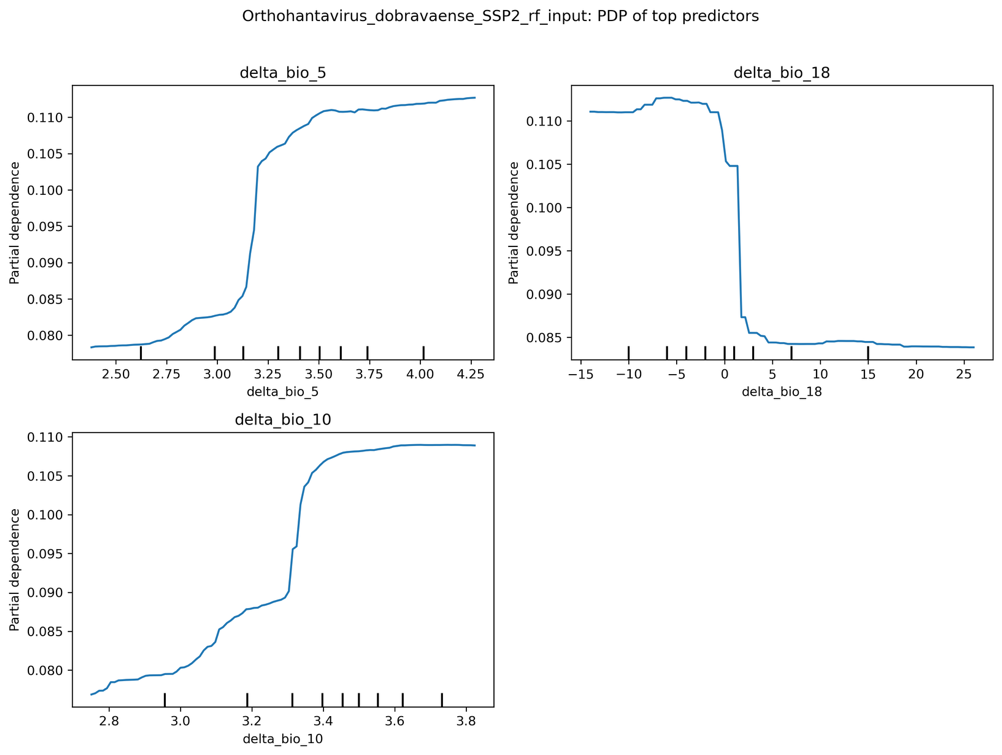

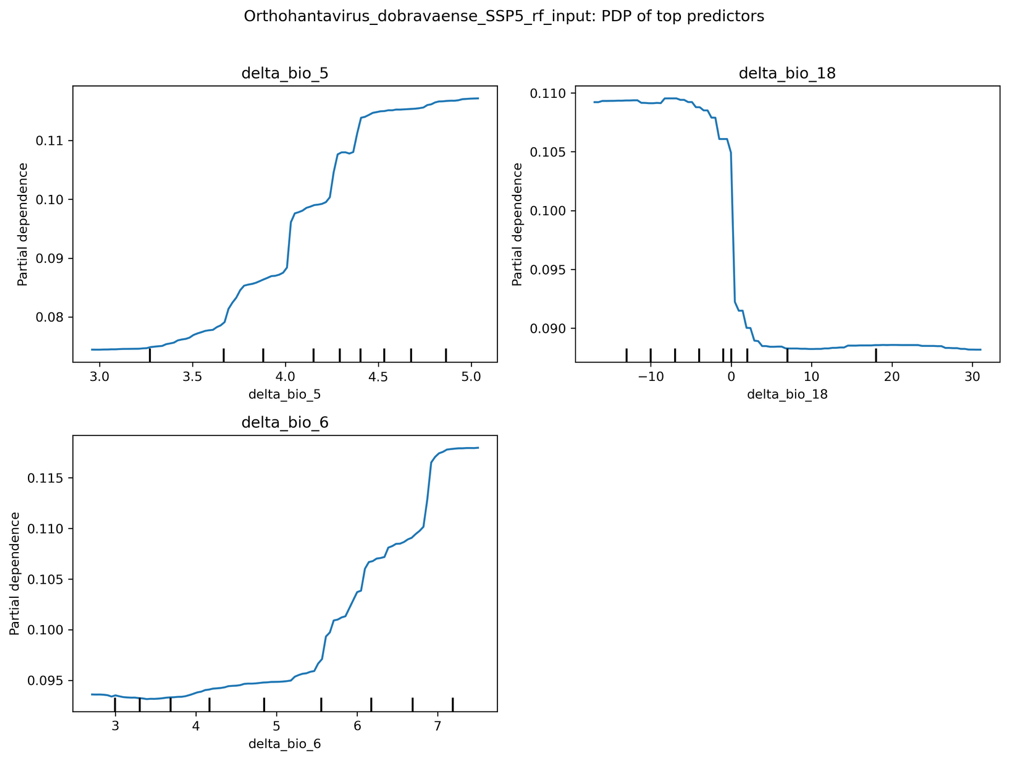

Fig S5.4. Orthohantavirus dobravaense

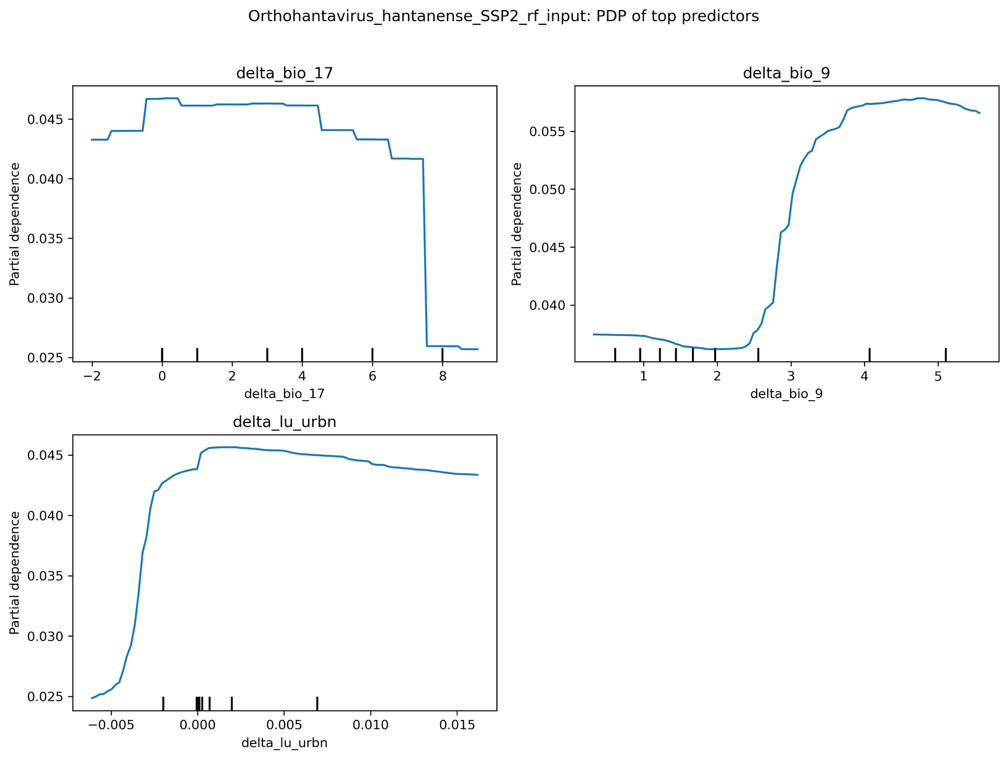

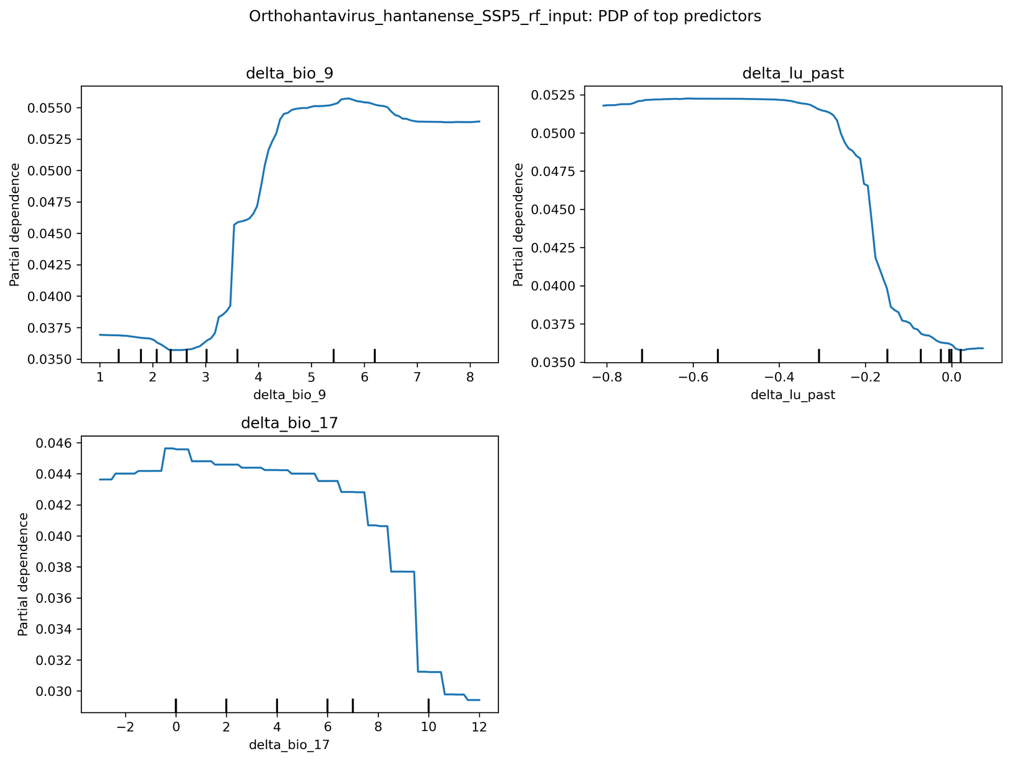

Fig S5.5. Orthohantavirus hantanense

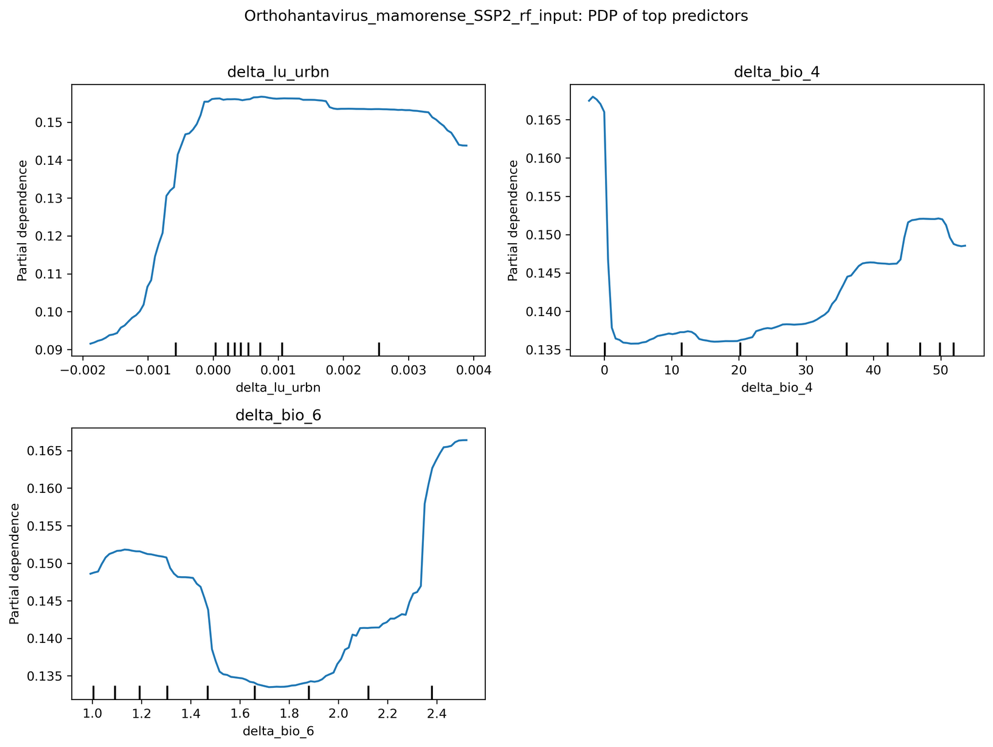

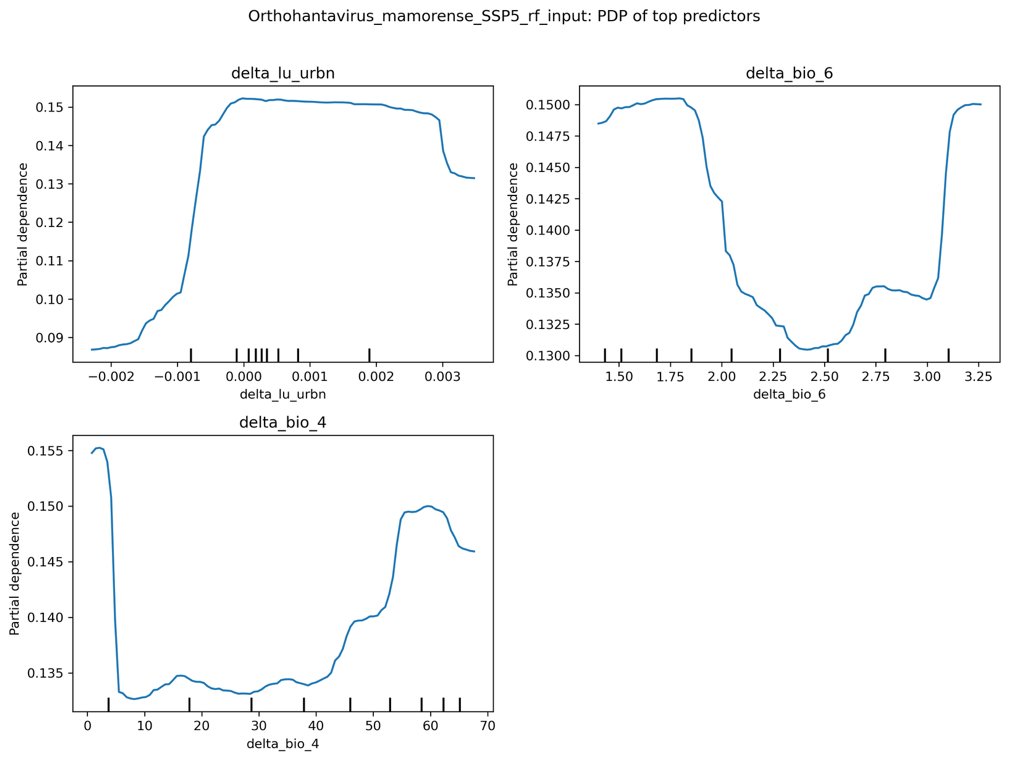

Fig S5.6. Orthohantavirus mamorense

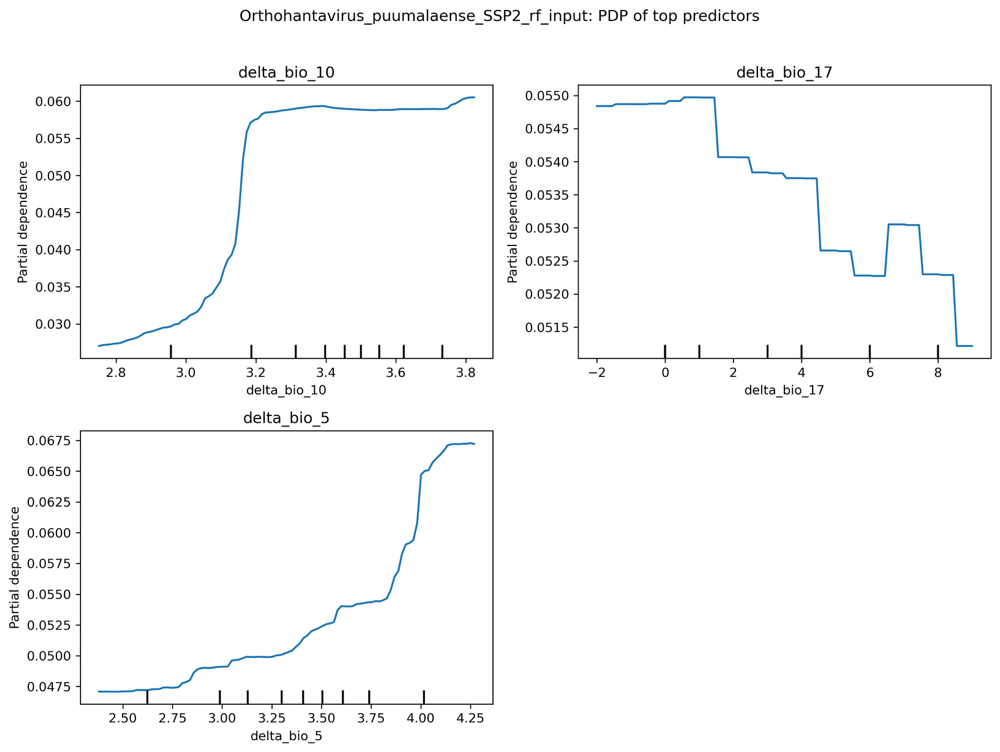

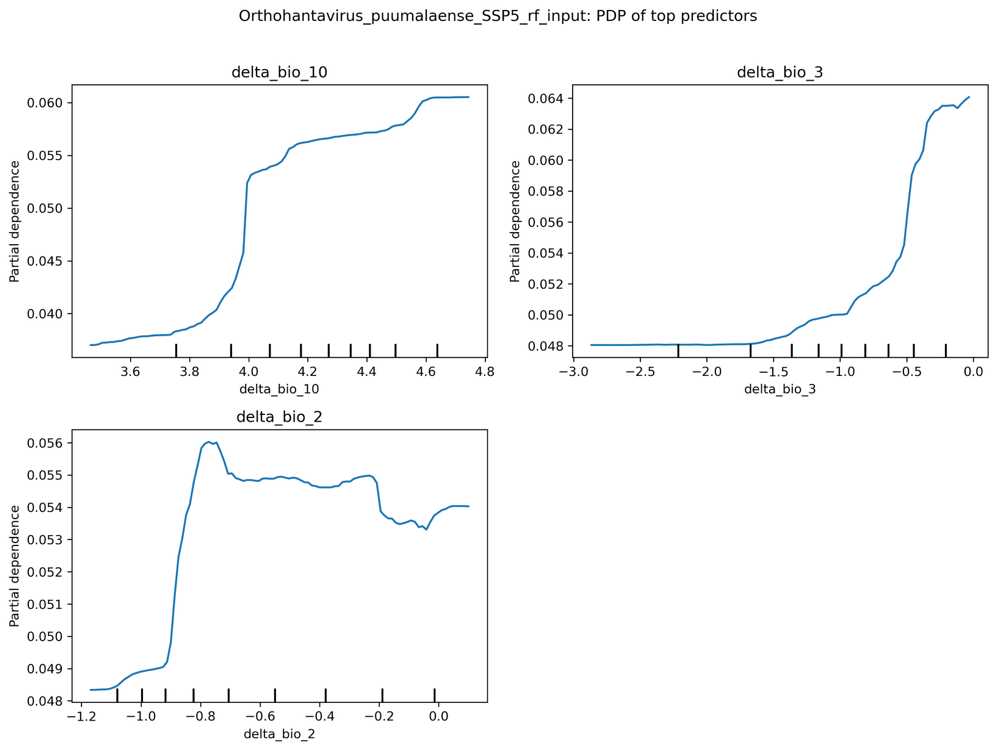

Fig S5.7. Orthohantavirus puumalense

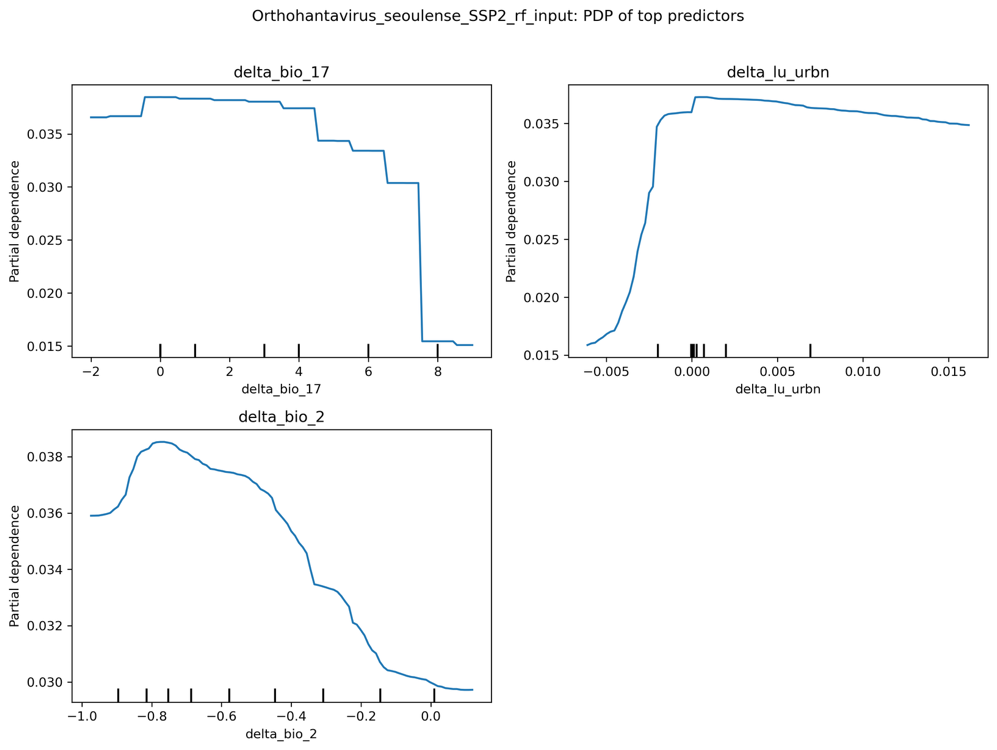

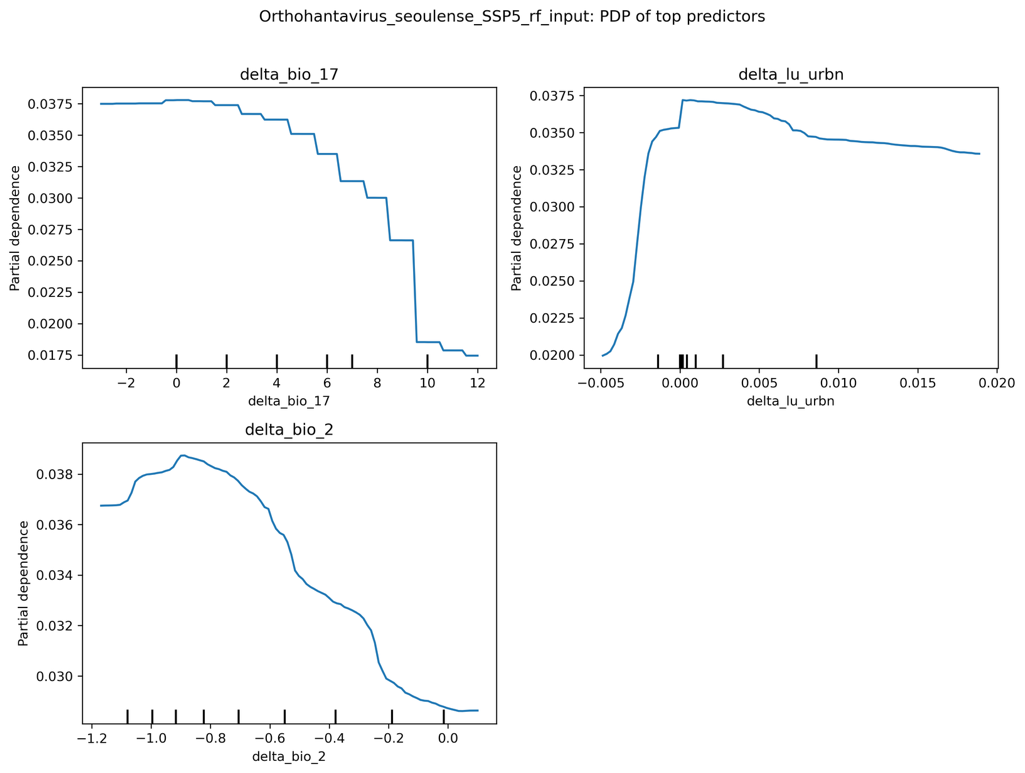

Fig S5.8. Orthohantavirus seoulense

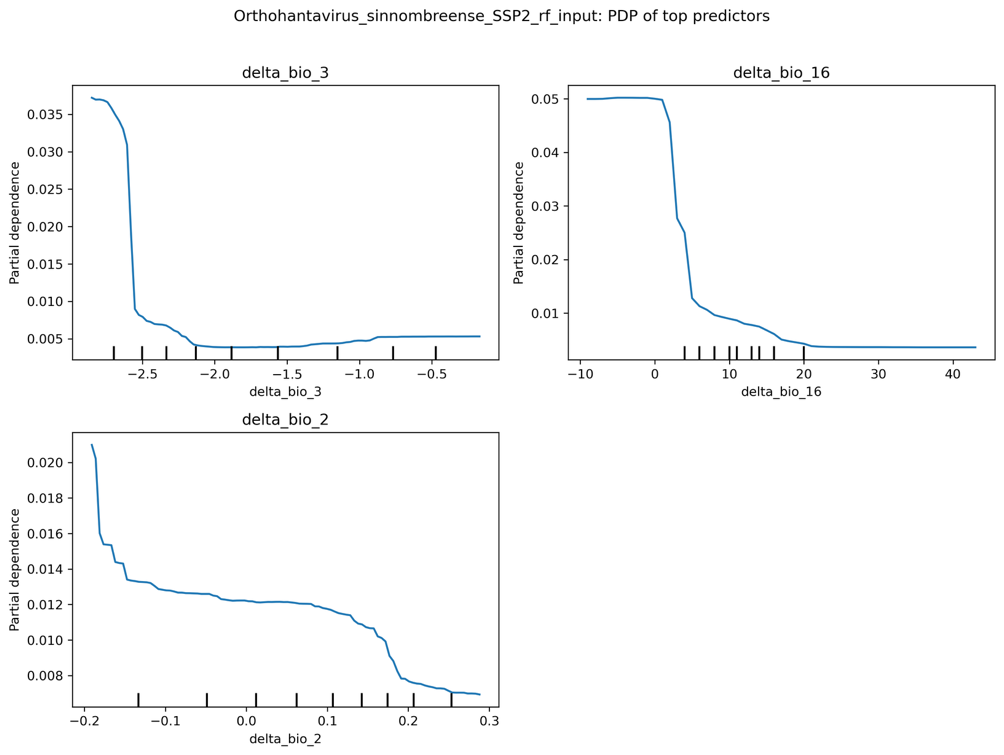

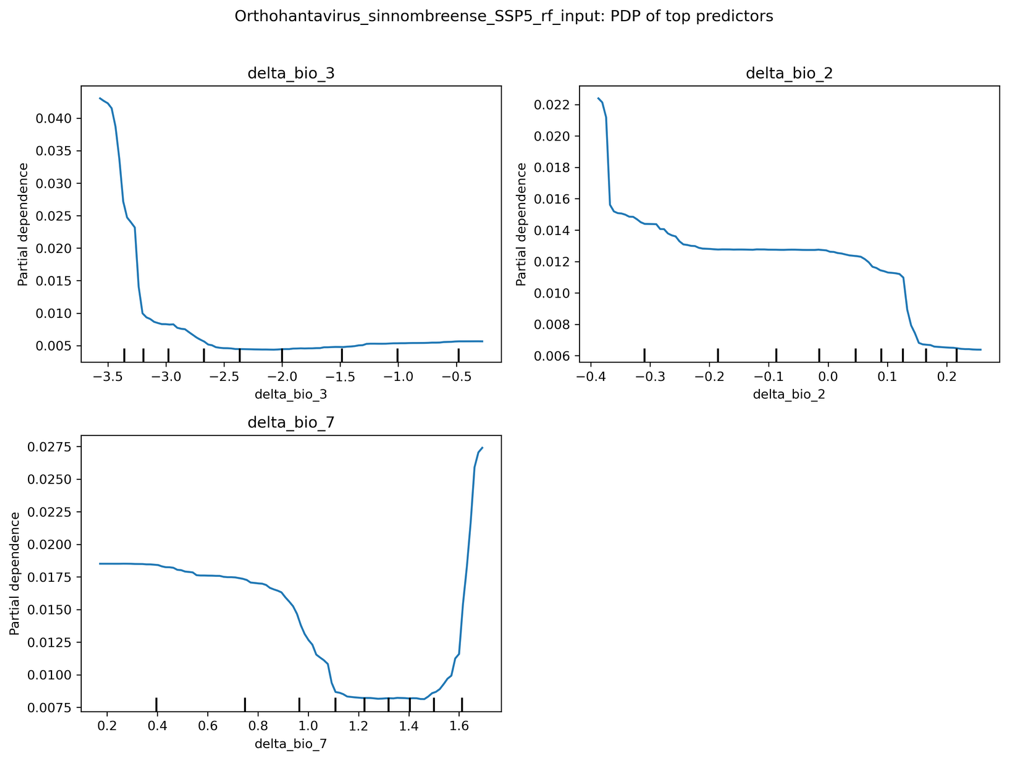

Fig S5.9. Orthohantavirus sinnombreense

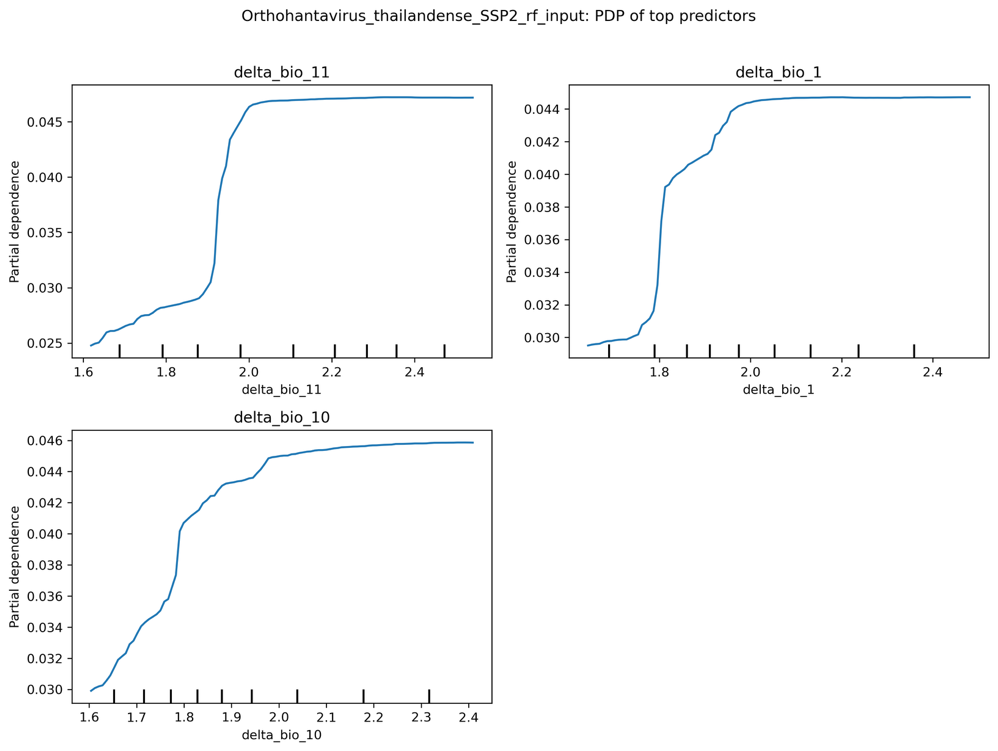

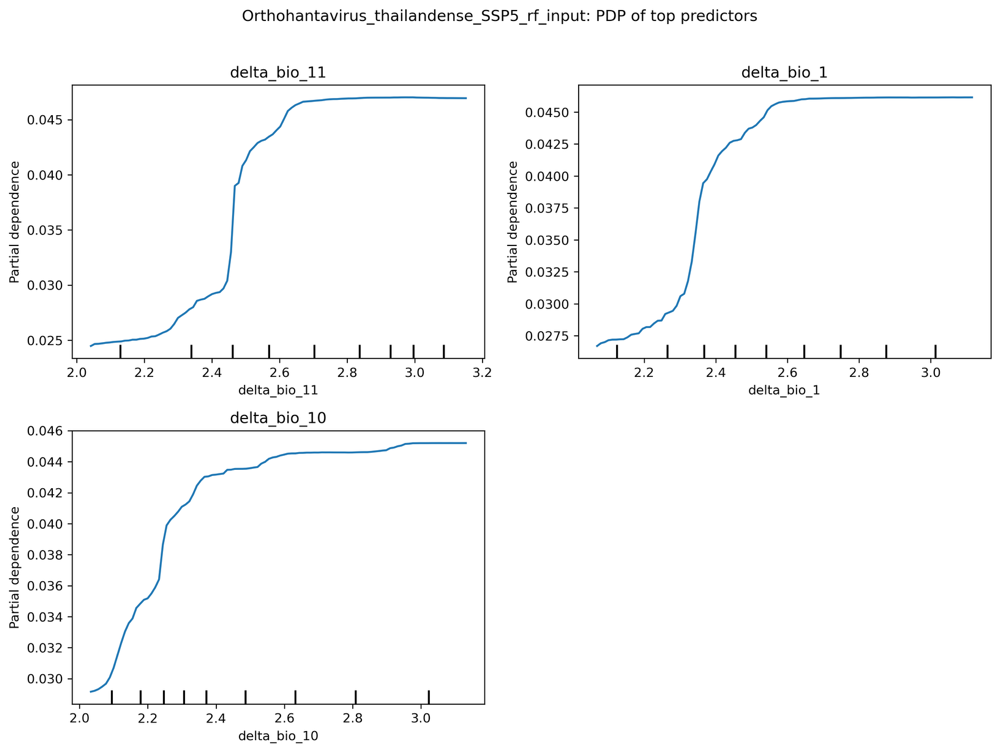

Fig S5.10. Orthohantavirus thailandense

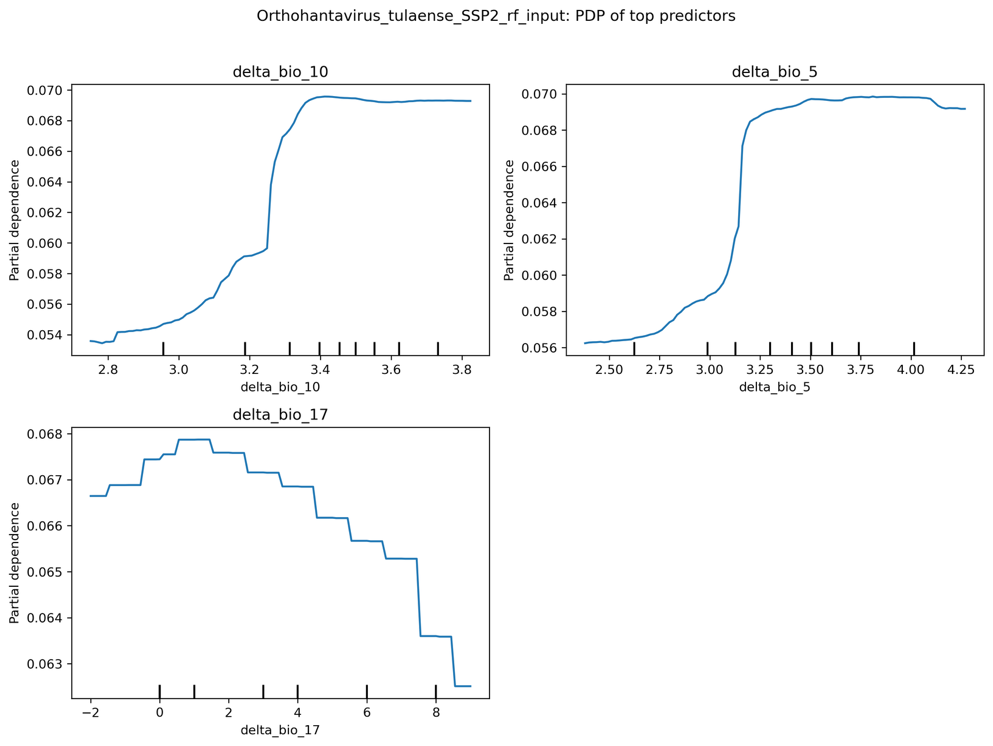

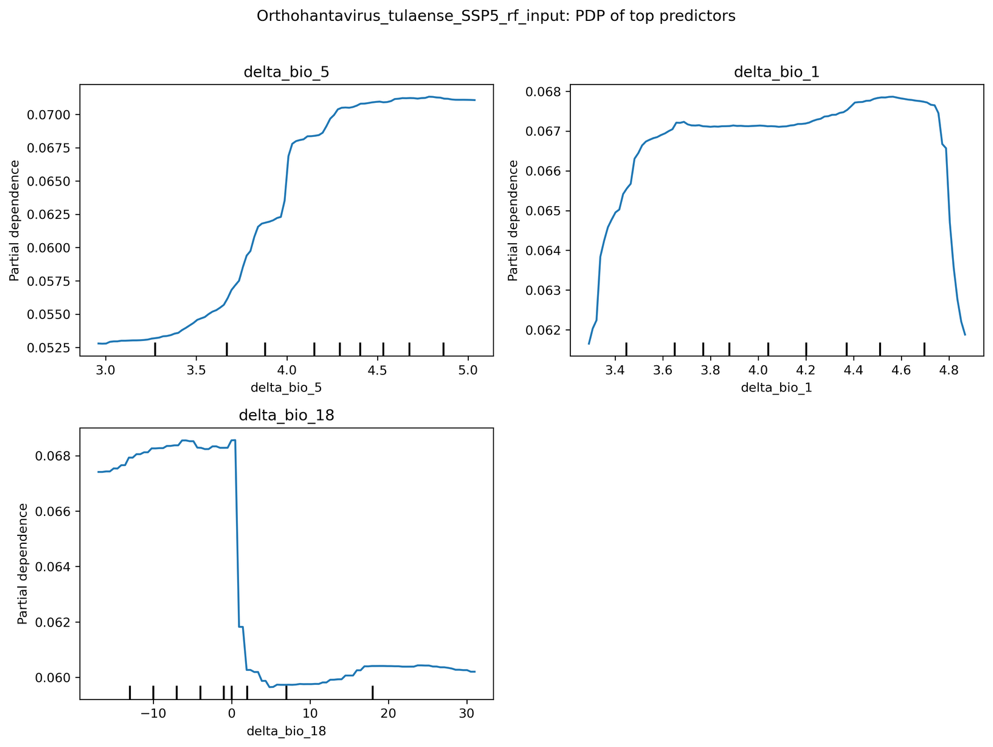

Fig S5.11. Orthohantavirus tulaense

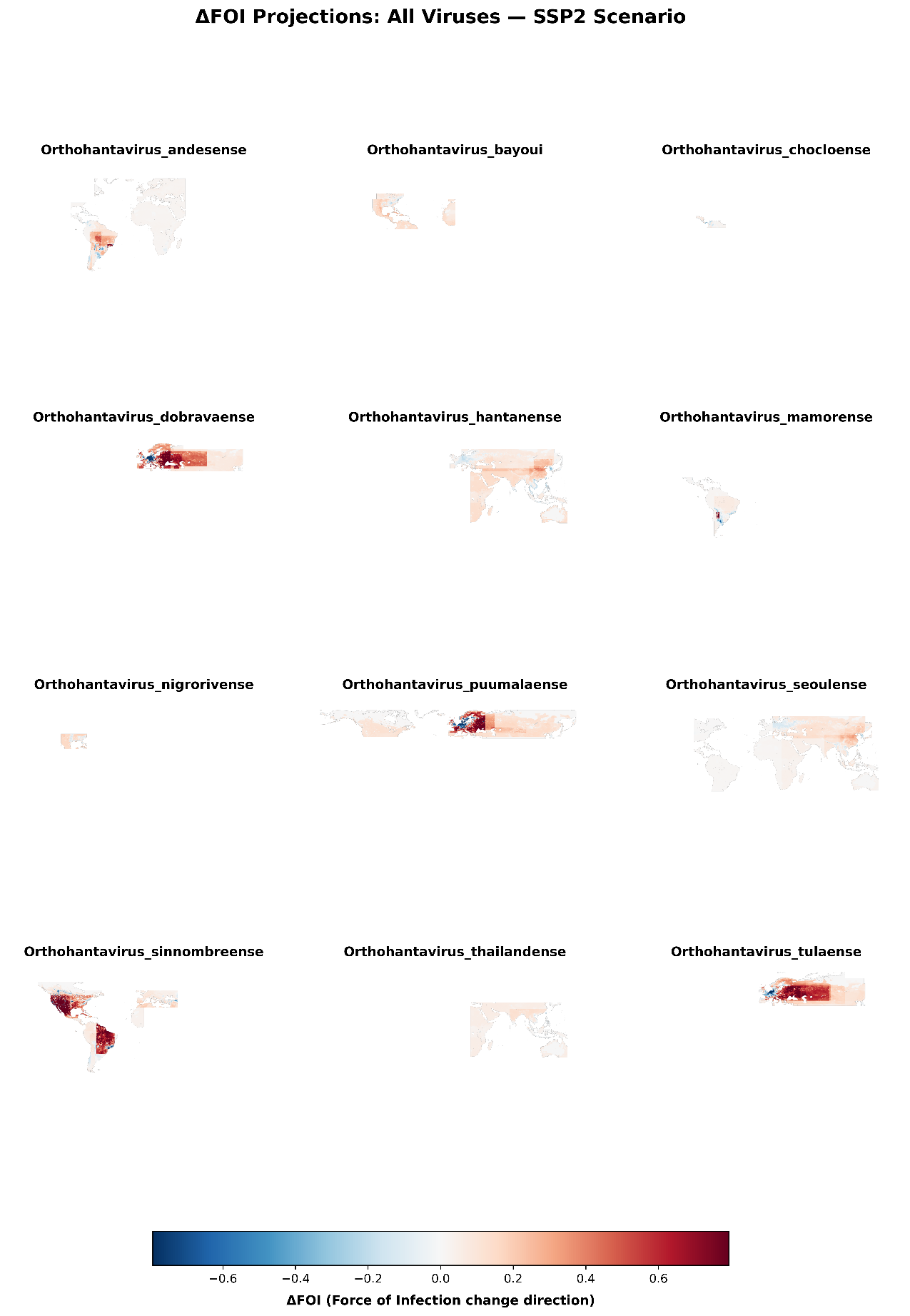

Fig S6 Virus specific climate-driven changes in spillover hazard. Differences in force-of-infection (FOI) estimates between current conditions and future projections under SSP2–4.5 illustrate how climate-mediated shifts in reservoir suitability reshape zoonotic risk.

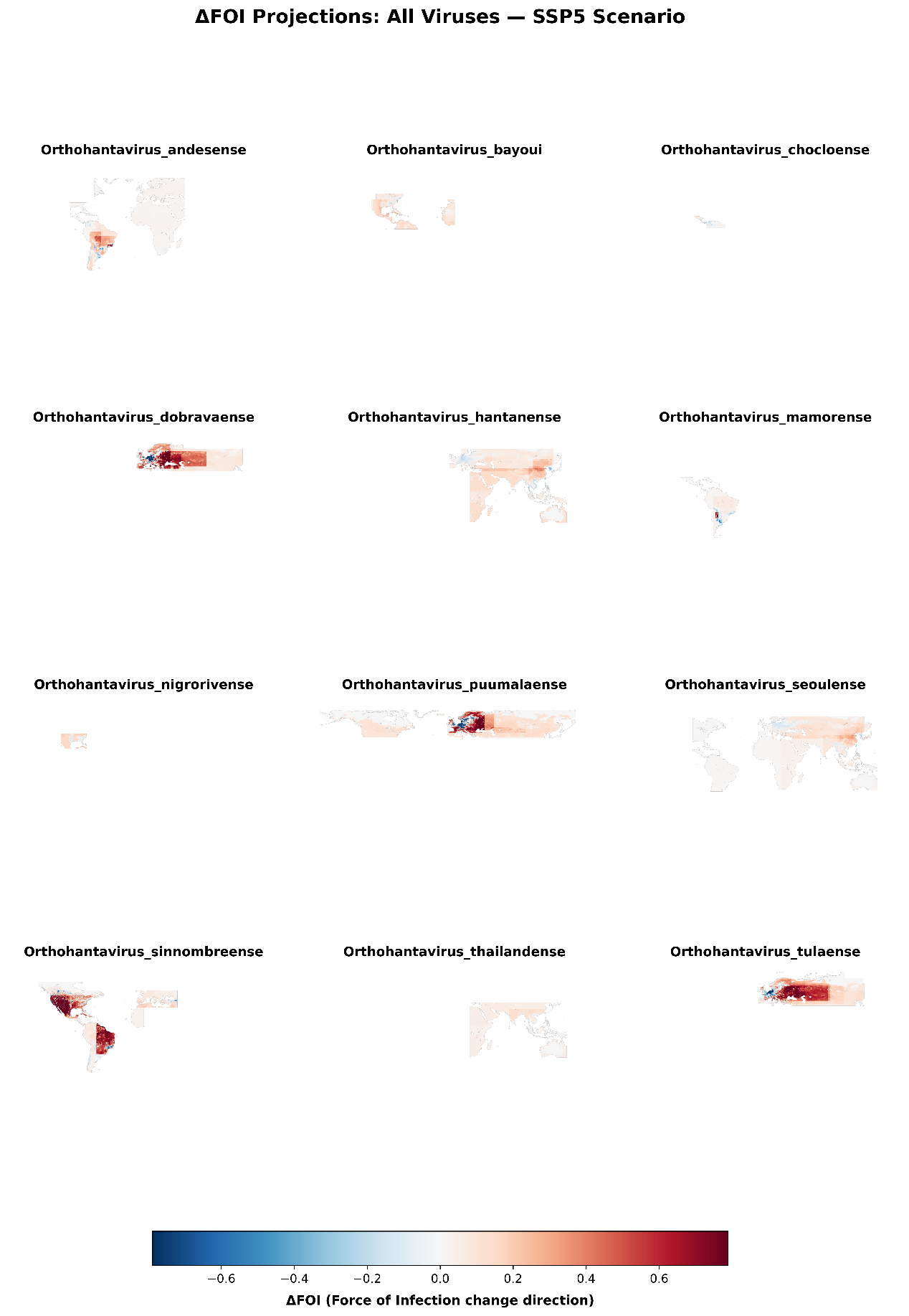

Fig S7 Virus specific climate-driven changes in spillover hazard. Differences in force-of-infection (FOI) estimates between current conditions and future projections under SSP5–8.5 illustrate how climate-mediated shifts in reservoir suitability reshape zoonotic risk.
